# LAP Drives Monocyte-Mediated and Extracellular Translocation of *Listeria monocytogenes* Across the Placental Barrier

**DOI:** 10.64898/2025.12.22.696054

**Authors:** Ayesha Atique, Breanna Amelunke, Zayd A. Hasan, Donald B. Bryant, Oliver R. Oakley, Marcia Pierce, Rishi Drolia

**Author notes:** **These authors contributed equally** (Ayesha Atique, Breanna Amelunke).

## Abstract

*Listeria monocytogenes* (*Lm*) is a major cause of fetal infection, yet the mechanisms by which it traverses the placental barrier remain incompletely defined. We established and validated a physiologically relevant in vitro human placental co-culture barrier model using BeWo trophoblasts and primary human placental vascular endothelial cells (HPVECs), which exhibited robust barrier integrity and organized tight junctions. Using this model, we demonstrate that *Lm* exploits a Trojan-horse transmigration mechanism via infected monocytes. *Lm*-infected THP-1 monocytes displayed a ∼4.8-fold higher monocyte transmigration and ∼8.12-fold greater intracellular bacterial delivery across the placental barrier compared with non-pathogenic *Listeria innocua (Li)*. Monocyte transmigration was associated with claudin-1 and occludin disruption, increased paracellular permeability, and localization of infected monocytes at junctional breach sites.

Monocyte-mediated placental traversal required the *Lm* virulence factors *Listeria* adhesion protein (LAP) and internalin B (InlB), as *lap⁻* and *ΔinlB* mutants exhibited *Li-*like transmigration and an ∼90% reduction in intracellular bacterial delivery, despite normal monocyte uptake. *Lm*-infected monocytes induced a ∼1.86-fold increase in sVCAM-1 secretion, which was LAP- and InlB-dependent, consistent with pro-transmigration endothelial phenotype.

In contrast, extracellular traversal required LAP and internalin A (InlA), as *lap⁻* and *ΔinlA* mutants exhibited severe defects comparable to *Li*, whereas *ΔinlB* and *Δhly* were minimally impaired. Exposure of monocytes to extracellular *Lm* further amplified transmigration and barrier permeability.

Together, these data define two distinct placental invasion routes: a monocyte-associated LAP/InlB-dependent pathway and an extracellular LAP/InlA-dependent pathway, identifying LAP as a critical noncanonical virulence factor in placental infection and mechanisms of vertical transmission.

## INTRODUCTION

*Listeria monocytogenes* (*Lm*) is a facultative intracellular foodborne pathogen capable of breaching the placental barrier, leading to severe fetal outcomes, including miscarriage, stillbirth, and neonatal infection (1–3). The placenta is a complex, multilayered barrier composed of syncytiotrophoblasts, cytotrophoblasts, and fetal endothelial cells, which collectively provide immune and physical protection to the fetus (4–7). Despite its robust defense, the placenta is uniquely vulnerable to *Lm* infection, driven in part by the pathogen’s specialized virulence factors (8, 9).

The ability of *Lm* to cross host barriers is facilitated by the bacterial surface proteins Internalin A (InlA) and Internalin B (InlB) (10, 11). InlA, covalently anchored to the bacterial cell wall, engages E-cadherin, a transmembrane adhesion molecule essential for maintaining epithelial adherens junctions (12). Internalin B (InlB), a noncovalently bound surface protein of *Lm*, targets the hepatocyte growth factor (HGF) receptor c-Met. By mimicking HGF signaling, InlB activates phosphoinositide 3-kinase (PI3-K), inducing membrane ruffling and facilitating bacterial entry (13). In transgenic mice expressing human E-cadherin, which is permissive to both InlA/B pathways, deletion of both *inlA* and *inlB* severely impacts fetal infection, highlighting their synergistic role in placental invasion (8, 14). In contrast, studies in the guinea pig model, where E-cadherin is inherently InlA-permissive, demonstrate that *inlA* deletion does not reduce placental barrier crossing. These findings indicate that direct InlA-mediated invasion of trophoblasts is dispensable *in vivo* and suggest that alternative virulence factors or mechanisms can predominate at the placental barrier. Moreover, while the roles of canonical internalins (InlA, InlB, and InlP) in placental barrier crossing are well established, non-canonical internalin-independent virulence factors contributing to *Lm* translocation across the placental barrier remain poorly defined (Disson et al., 2008; Faralla et al., 2018). Importantly, clinical strains with truncated or absent forms of the canonical internalins can still cause severe neonatal infections in humans, guinea pigs, and mice (Bakardjiev et al., 2004; Cruz et al., 2013; Fravalo et al., 2017; Gelbíčová et al., 2015; Ghanem et al., 2012; Oliver et al., 2013), strongly implicating internalin-independent pathways in placental barrier translocation.

*Listeria* adhesion protein (LAP), a 94-kDa alcohol acetaldehyde dehydrogenase (AdhE *lmo1634*), is a SecA2-dependent surface-associated and secreted moonlighting protein (1, 15). LAP promotes paracellular translocation across the intestinal epithelial barrier by binding to its cognate host receptor, heat shock protein 60 (Hsp60)(16, 17). LAP-Hsp60 engagement activates NF-κB signalling and disrupts tight junctions via myosin light chain kinase (MLCK)-mediated and caveolin-1-dependent remodelling (1, 17–19). However, LAP’s role in traversing the placental barrier remains unexplored. Listeriolysin O (LLO), encoded by the *hly* gene, while essential for phagosomal escape, does not directly mediate barrier crossing (20, 21).

Beyond direct cellular invasion by the extracellular blood-borne bacteria, as a facultative intracellular pathogen, *Lm* may exploit a Trojan-horse mechanism, whereby infected monocytes or dendritic cells facilitate bacterial dissemination across host barriers (22, 23). In the context of neuroinvasion, *Lm* associates with monocytes, enabling lateral transfer of bacteria across the blood-brain barrier via cell-to-cell spread, without clear evidence that the monocytes themselves transmigrate (24). InlB contributes to this process by engaging c-Met–mediated PI3K/Akt signalling to promote intracellular survival and by suppressing apoptosis through FLICE-like inhibitory protein (FLIP)-dependent inhibition of caspase-8 activation (24). While these findings suggest that *Lm* exploits monocytes to gain barrier access, whether infected monocytes can themselves actively transmigrate across barriers, such as the placenta, and deliver viable intracellular bacteria remains unknown. Furthermore, the extent to which monocyte transmigration contributes to vertical transmission has not been systematically investigated in a physiologically relevant human model.

Experimental models investigating *Lm* translocation have traditionally relied on murine systems or single-cell invasion assays, which do not fully recapitulate the complex trophoblast–endothelial architecture of the human placenta. Mouse and guinea pig models are commonly used to study infection; however, species-specific differences limit their translational relevance. In particular, the InlA–E-cadherin interaction is not conserved in mice, and InlB fails to engage c-Met in guinea pigs, while both models differ from humans in placental organization (dichorial vs. monochorial) (25, 26). Recent co-culture systems offer improved modeling of human placental barrier properties (27–29). Consistent with a broader shift in NIH priorities, there is a growing emphasis on human-relevant, non-animal platforms to better model host–pathogen interactions and reduce reliance on animal systems (30).

To address these gaps, we developed and validated a human placental co-culture barrier model and demonstrate that *Lm* exploits a Trojan-horse mechanism in which *Lm*-infected monocytes show markedly enhanced transmigration, disrupt claudin-1 and occludin junctions, and deliver intracellular bacteria across the barrier. This monocyte-mediated traversal requires LAP and InlB, which drive monocyte migration and induce endothelial sVCAM-1 upregulation. In parallel, extracellular *Lm* crossing requires LAP and InlA, with LAP emerging as the shared, essential determinant across both pathways. Collectively, these findings identify LAP as a critical virulence factor enabling both monocyte-associated and extracellular modes of placental barrier traversal.

## RESULTS

### Establishment and Permeability Characterization of a Human Placental Co-culture Model Using BeWo and HPVECs

To reconstruct the physiologically relevant placental barrier, we developed and validated an in vitro co-culture Transwell model in which BeWo trophoblasts and primary human placental vascular endothelial cells (HPVECs) were seeded on the apical and basal surfaces of permeable inserts, respectively (**Fig. 1A**). Phase-contrast microscopy of the co-culture model at 72 h post-seeding showed formation of stable, confluent monolayers (**Fig. 1B**).

**FIG 1.**
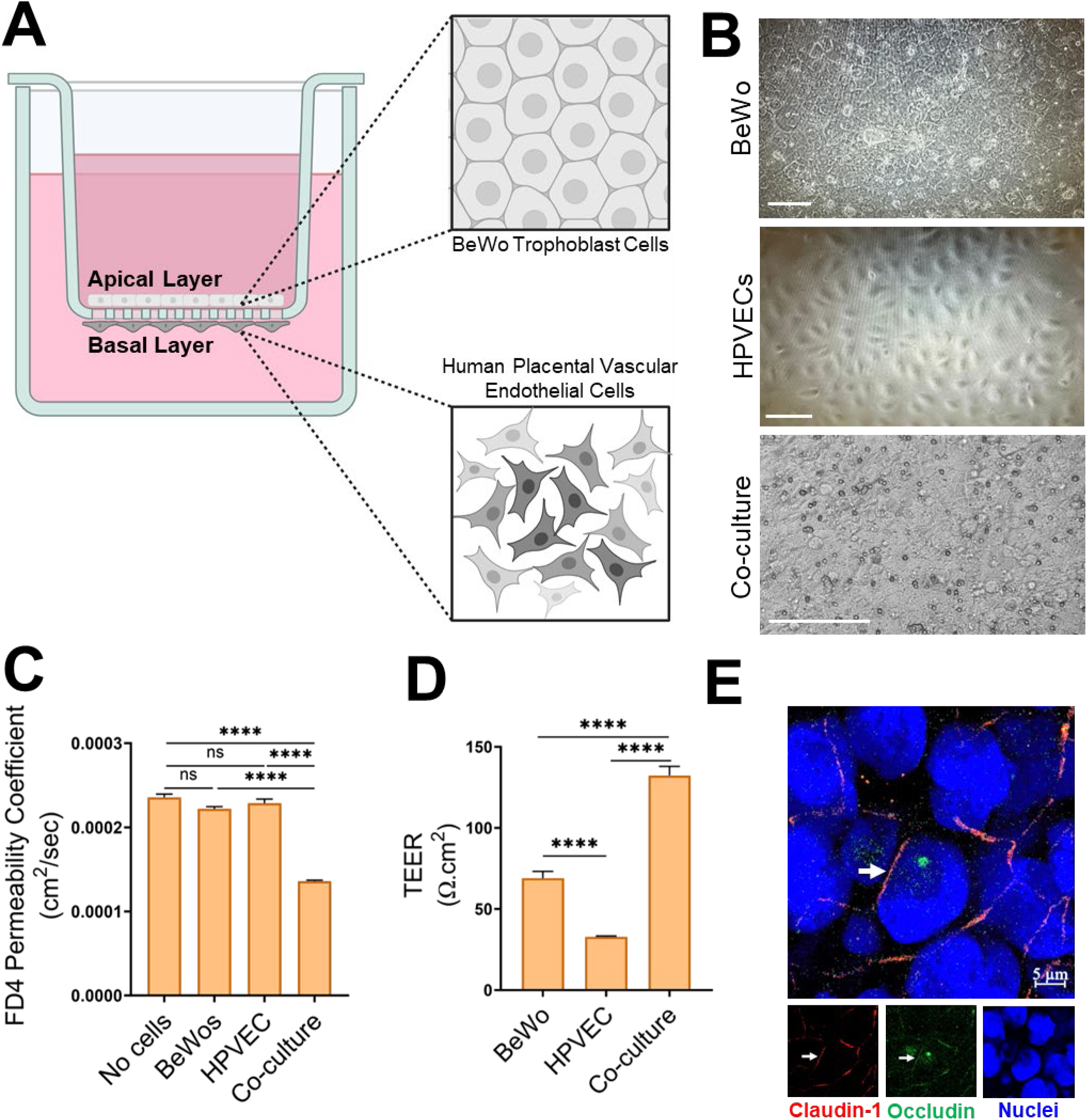
Validation of a dual-layered human placental co-culture model. (**A**) Schematic illustration of the co-culture model with BeWo cytotrophoblast-like cells seeded on the apical surface and primary human placental vascular endothelial cells (HPVECs) seeded on the basal surface of Transwell inserts. (**B**) Phase-contrast images of BeWo (top) and HPVECs (middle), cultured under standard conditions (scale bars, 200µm). Phase-contrast images of the co-culture model (bottom) with BeWo and HPVECs seeded on opposite sides of a Transwell insert to recapitulate the placental barrier 72 h post-seeding (scale bars, 200µm). (**C, D**) FITC–dextran (4 kDa; FD4) permeability coefficient (C; n = 3 per condition) and Trans-epithelial electrical resistance (TEER; D; n =11 per condition) assessed at 72 h post-seeding, demonstrating enhanced barrier integrity in the co-culture relative to BeWo or HPVECs monolayers. Data represent mean ± SEM from at least three independent experiments. Statistical significance was determined by one-way ANOVA with Tukey’s multiple-comparisons test. ns, not significant; ****, P < 0.0001. (**E**) Representative confocal fluorescence micrograph of a co-culture model at 72 h post-seeding following immunostaining for claudin-1 (red) and occludin (green). Nuclei were counterstained with DAPI (blue). Individual fluorescence channels are shown below the merged image.

At this timepoint, a fluorescein isothiocyanate (FITC)–dextran 4 kDa (FD4) permeability assay demonstrated that the co-culture model possessed the lowest barrier permeability, exhibiting a 38.7% reduction relative to BeWo monocultures and a 40.5% reduction compared with HPVECs monocultures (**Fig. 1C**). Consistently, transepithelial electrical resistance (TEER) measurements indicated markedly enhanced barrier function, with the co-culture displaying a 91.7% increase in resistance compared with BeWo alone and a 302% increase relative to HPVECs (**Fig. 1D**).

Immunofluorescence analysis further confirmed the presence of continuous claudin-1 and occludin tight junction localization across the co-culture barrier (**Fig. 1E**). Together, these results validate the structural and functional integrity of the dual-layered placental model and support its use for downstream permeability and transmigration studies.

### Infected THP-1 Monocytes Mediate Transmigration of *Lm* Across the Placental Co-culture Model

We first quantified the intracellular uptake of wild-type (WT) *Lm* and the non-pathogenic species *Listeria innocua* (*Li*) in THP-1 monocytes. At one hour post-infection (hpi) followed by 30 minutes of gentamicin treatment to eliminate extracellular bacteria, intracellular colony-forming units (CFUs) were enumerated. *Li*, which lacks the *Listeria* pathogenicity island-1 (LIPI-1), achieved early intracellular burdens comparable to those of WT *Lm* (**Fig. 2A**). Giemsa-stained Cytospin preparations confirmed the presence of gentamicin-protected intracellular bacteria in both *Lm-* and *Li-*infected THP-1 cells (**Fig. 2B**). Quantitative analysis of gentamicin-protected intracellular bacterial burdens revealed no significant difference between the two strains, indicating similar monocyte uptake (**Fig. 2C**). Extending gentamicin exposure by an additional 2.5h after 1hpi also yielded similar intracellular CFU burdens, indicating no early survival defect in *Li* (**Fig. 2A**).

**FIG 2.**
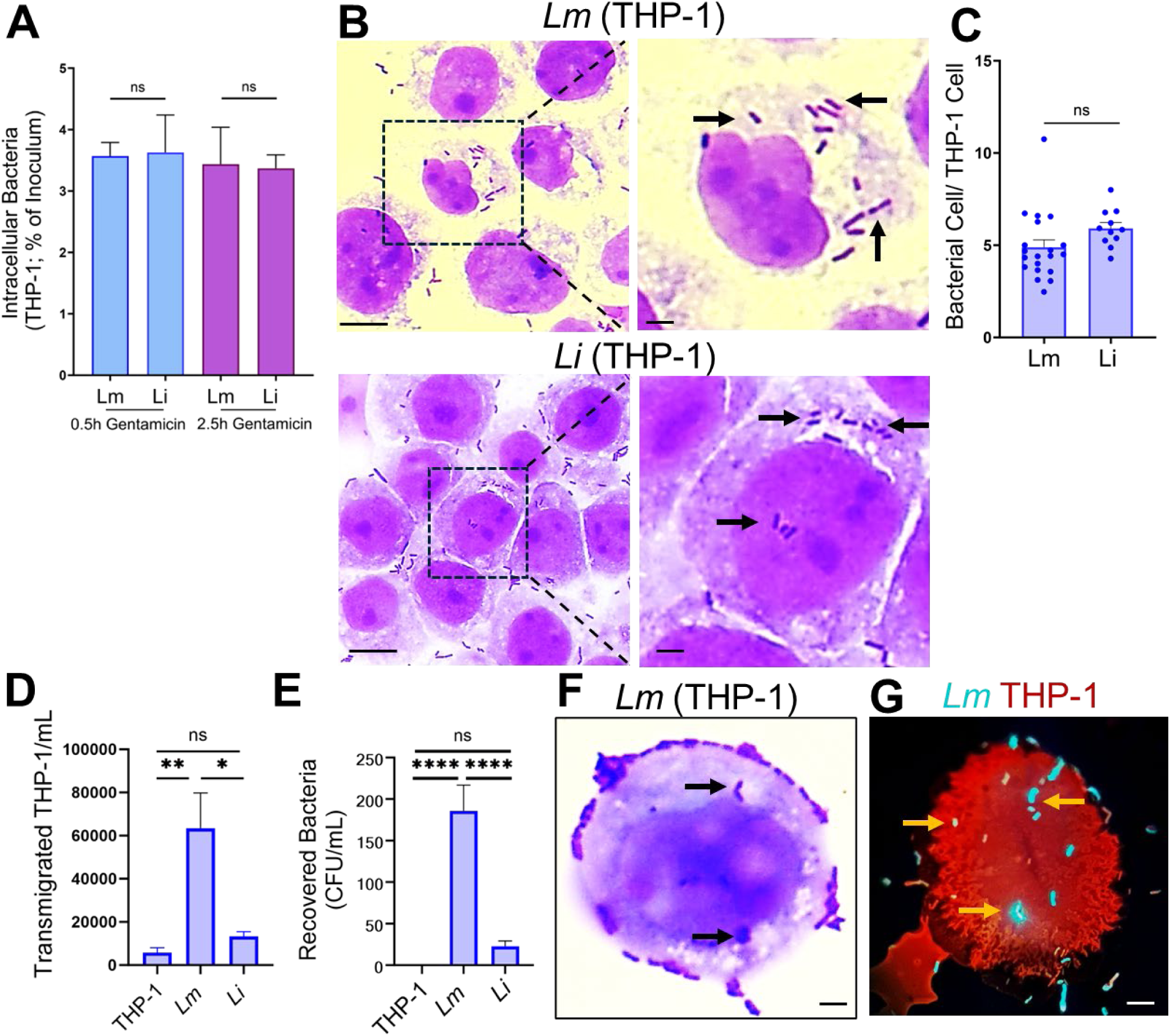
Transmigration of *Listeria monocytogenes* via infected THP-1 monocytes across the placental co-culture model. (**A–C**) Intracellular burden and visualization of *L. monocytogenes* (*Lm*) and *L. innocua* (*Li*) in THP-1 monocytes. THP-1 cells (1×10⁶) were infected with *Lm* or *Li* (MOI 50, 1 h) and treated with gentamicin sulfate (50 µg/mL) for 30 min or 2.5 h to eliminate extracellular bacteria. (**A**) Intracellular bacteria were quantified post-lysis and expressed as the percentage of the initial inoculum recovered. *Lm* and *Li* showed no significant difference in intracellular burden at both time points. (n = 3 per treatment) (**B**) Representative Giemsa-stained Cytospins of gentamicin-protected THP-1 cells infected with *Lm* or *Li* (MOI 50, 1 h + 30 min gentamicin sulfate). Scale bar, 20 µm. Insets (right) highlight intracellular bacteria (arrows). Inset scale bar, 1 µm. (**C**) Quantification of intracellular bacteria per THP-1 cell from Giemsa-stained images (dots represent the average of 10–15 cells/condition; n = 150–175 cells per treatment). (**D**) Quantification of transmigrated THP-1 monocytes recovered in the basolateral chamber of the co-culture placental model after 2 h transmigration. n = 6-8 per treatment (**E**) Quantification of intracellular bacteria recovered from transmigrated THP-1 from the basolateral chamber of the co-culture placental model after 2 h transmigration. n = 7 per treatment (**F, G**) Representative Cytospin micrographs of gentamicin-protected THP-1 cells recovered from the basolateral chamber showing intracellular *Lm* (arrows), visualized by Giemsa staining (**F**) or CellTracker Red–labeled THP-1 cells containing GFP-labeled *Lm* (**G**). Scale bar, 1 µm. Data in A, C, D, and E represent the mean ± SEM from at least three independent experiments. Statistical significance was determined by one-way ANOVA with Tukey’s multiple-comparisons test. ns, not significant; *, P < 0.05; **, P < 0.01; ****, P < 0.0001.

Given the comparable early intracellular burdens between *Lm* and *Li*, we next assessed whether infected monocytes differ in their ability to transmigrate across the placental co-culture model. THP-1 monocytes, either uninfected or infected with WT *Lm* or *Li*, were added to the apical chamber of a Transwell insert containing the dual-layered model. During the 2-hour transmigration period, both apical and basolateral chambers were treated with gentamicin to eliminate extracellular bacteria. Migrated THP-1 cells were then collected from the basolateral compartment for analysis.

Quantification revealed a 4.8-fold increase in transmigration by *Lm*-infected THP-1 monocytes compared with *Li*-infected monocytes (**Fig. 2D**). Consistent with this, intracellular *Lm* CFUs recovered from the basolateral THP-1 were 8.12-fold higher than those of *Li* (**Fig. 2E**), indicating more efficient delivery of viable intracellular bacteria across the barrier by *Lm*-infected monocytes. Giemsa-stained Cytospin preparations of the basolateral samples confirmed the presence of gentamicin-protected intracellular *Lm* within the transmigrated THP-1 cells (**Fig. 2F)**.

To directly visualize intracellular carriage of bacteria during transmigration, we used GFP-expressing *Lm* and CellTracker Red–labeled THP-1 cells (**Fig. S1A-D)**. Following the transmigration assay, GFP-positive *Lm* were observed within CellTracker Red–labeled THP-1 cells in Cytospin preparations of media collected from the basolateral compartment (**Fig. 2G)**. Together, these data show that *Lm*-infected monocytes exhibit enhanced transmigration and deliver higher numbers of viable intracellular bacteria across the placental barrier co-culture model.

### *Lm*-Infected Monocytes Compromise Placental Barrier Tight Junction Integrity to Facilitate Transmigration

Tight junctions at the placental barrier regulate monocyte transmigration and can limit monocyte–mediated entry of pathogens (31). To assess whether *Lm*-infected monocytes compromise barrier function, we examined tight junction integrity and permeability. Exposure to uninfected THP-1 monocytes caused minor disruptions, with occludin and claudin-1 staining largely resembling intact junctions (**Fig. 3A**). In contrast, *Lm*-infected monocytes induced pronounced fragmentation and loss of both claudin-1 and occludin (**Fig. 3A**). Quantification confirmed minimal disruption in controls (claudin-1: 4.7%; occludin: 7.3%), moderate disruption with uninfected THP-1 cells (claudin-1: 15.0%; occludin: 18.3%), and marked disruption with *Lm*-infected monocytes (claudin-1: 72.3%; occludin: 79.0%) (**Fig. S2**).

**FIG 3.**
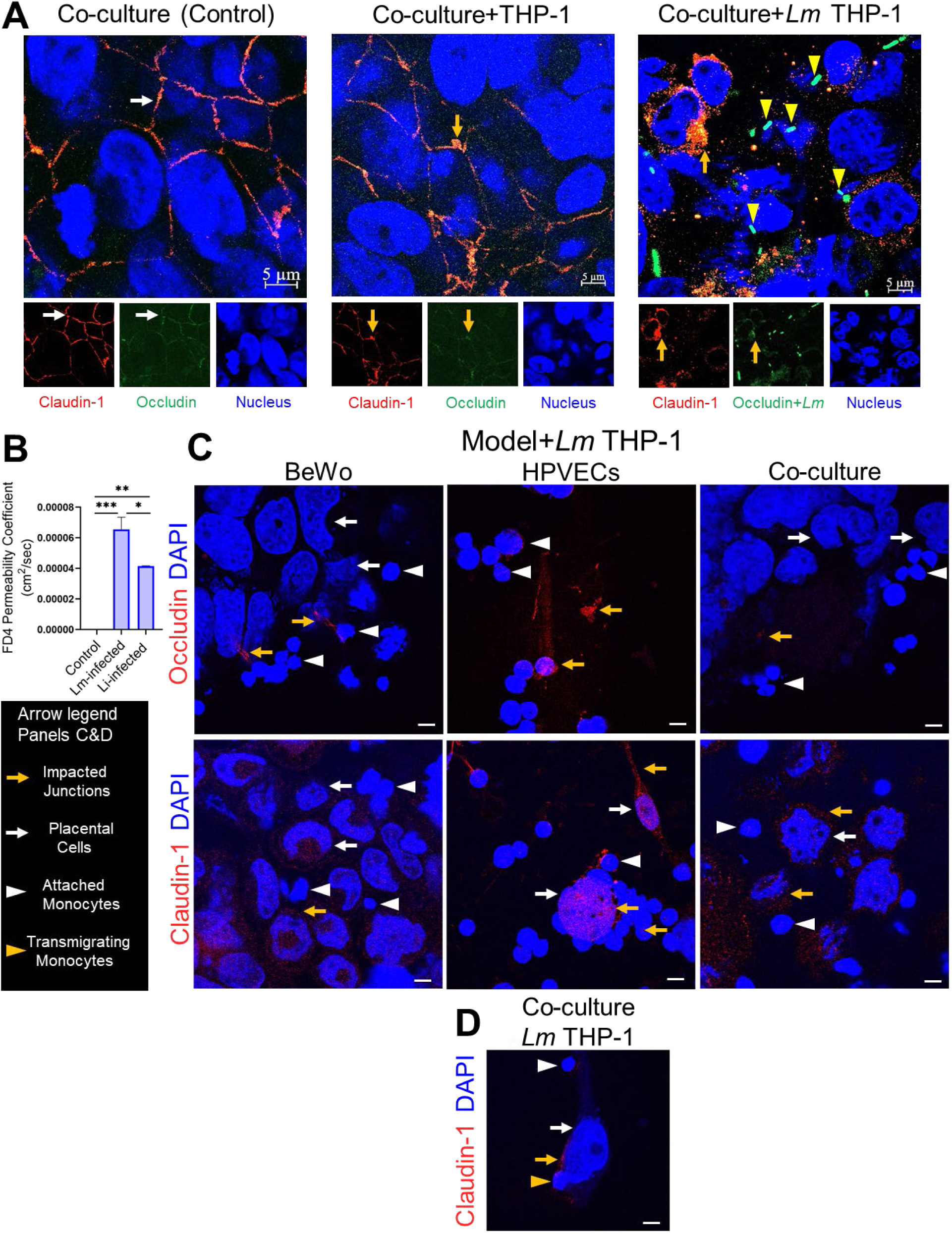
*Lm*–infected monocytes disrupt placental barrier junctions and increase permeability. (**A**) Representative confocal fluorescence micrographs of placental co-culture controls: untreated (left), exposed to uninfected THP-1 monocytes (middle), or exposed to *Listeria monocytogenes (Lm)*–infected THP-1 monocytes (right) for 2 h. White arrows indicate intact junctions in untreated controls. Yellow arrows indicate minor claudin-1 and occludin perturbations in co-cultures exposed to uninfected THP-1 cells, whereas pronounced claudin-1 and occludin disruption is observed following exposure to *Lm*-infected THP-1 monocytes. *Lm-*GFP is shown in green. Yellow arrowheads denote gentamicin-protected, cell-associated *Lm.* Nuclei were stained with DAPI (blue), claudin-1 with Alexa Fluor 555 (red), and occludin with Alexa Fluor 488 (green). Individual fluorescence channels are shown below the merged images. Scale bars, 5 µm. (**B**) Placental barrier permeability to FITC–dextran (4 kDa; FD4) following apical exposure to *Lm*- or *Listeria innocua* (*Li*)–infected THP-1 monocytes for 2 h. FD4 fluorescence in basolateral media revealed significantly increased permeability in co-cultures exposed to *Lm*-infected monocytes compared with *Li*-infected monocytes. n = 3 per treatment. Statistical significance was determined by one-way ANOVA with Tukey’s multiple-comparisons test. Data represent mean ± SEM from three independent experiments. *, P < 0.05; **, P < 0.01; ***, P < 0.001 (**C**) Representative confocal micrographs of occludin (top) and claudin-1 (bottom) junctions in BeWo monocultures, HPVECs monocultures, and BeWo–HPVECs co-cultures exposed to *Lm*-infected THP-1 monocytes for 2 h. DAPI (blue) staining enables the distinction of monocytes from placental cells based on nuclear morphology. Scale bars, 5 µm. The left panel provides arrow key definitions for panels (C) and (D): yellow arrows indicate disrupted junctions, white arrows denote placental cells, white arrowheads indicate attached monocytes, and yellow arrowheads denote transmigrating monocytes. (**D**) Representative confocal micrographs of claudin-1 junctions in BeWo–HPVECs co-cultures during THP-1 transmigration. DAPI (blue) and claudin-1 (red) staining distinguish monocytes from placental cells. Yellow arrowheads indicate transmigrating monocytes, white arrows denote placental cells, and yellow arrows mark disrupted junctions. Scale bars, 5 µm.

Intracellular gentamicin-protected *Lm* was observed with infected cells occasionally engaging and contacting underlying placental cells (**Fig. 3A**). Consistent with these structural defects, FD4 permeability increased by 58% after exposure to *Lm*-infected monocytes versus *Li*-infected monocytes (**Fig. 3B**), indicating *Lm*-specific barrier compromise.

Immunofluorescence in both monoculture and co-culture revealed direct interactions between infected monocytes and tight junction complexes, with monocytes readily distinguished from placental cells by nuclear morphology (**Fig. 3C**; occludin, top panel; claudin-1, bottom panel). Confocal imaging in the co-culture model further demonstrated that transmigrating monocytes localized at sites of tight-junction discontinuity, supporting a mechanistic role for junctional remodeling in monocyte-assisted barrier crossing (**Fig. 3D**).

Together, these results indicate that *Lm*-infected monocytes actively disrupt placental tight junctions and increase paracellular permeability, thereby facilitating monocyte–mediated bacterial translocation across the placental barrier.

### LAP and InlA Promote Extracellular *Lm* Translocation Across the Co-culture Placental Model

Direct extracellular invasion of the placental barrier by canonical internalins (InlA, InlB) has been reported (8, 14), but the contributions of internalin-independent virulence factors at the placental barrier remain unclear. Using a trophoblast–endothelial co-culture Transwell model, we quantified translocation of WT F4244 and its isogenic mutants (Δ*inlA*, Δ*inlB*, *lap*⁻), as well as WT 10403S and Δ*hly* (LLO), added apically. Basolateral CFUs were enumerated at 2 hpi.

The *lap⁻* and *ΔinlA* mutants exhibited substantial defects in translocation, with 74.7% and 94.1% decreases relative to WT F4244, respectively, whereas Δ*inlB* showed WT-like levels and Δ*hly* displayed only a modest, non-significant reduction (**Fig. 4A & B**). Translocation deficiencies of *lap⁻* and *ΔinlA* mutants were comparable to *Li*. Consistently, WT *Lm* induced a 180% increase in FD4 permeability relative to *Li*, indicating greater barrier disruption by extracellularly invading *Lm* (**Fig. 4C**).

**FIG 4.**
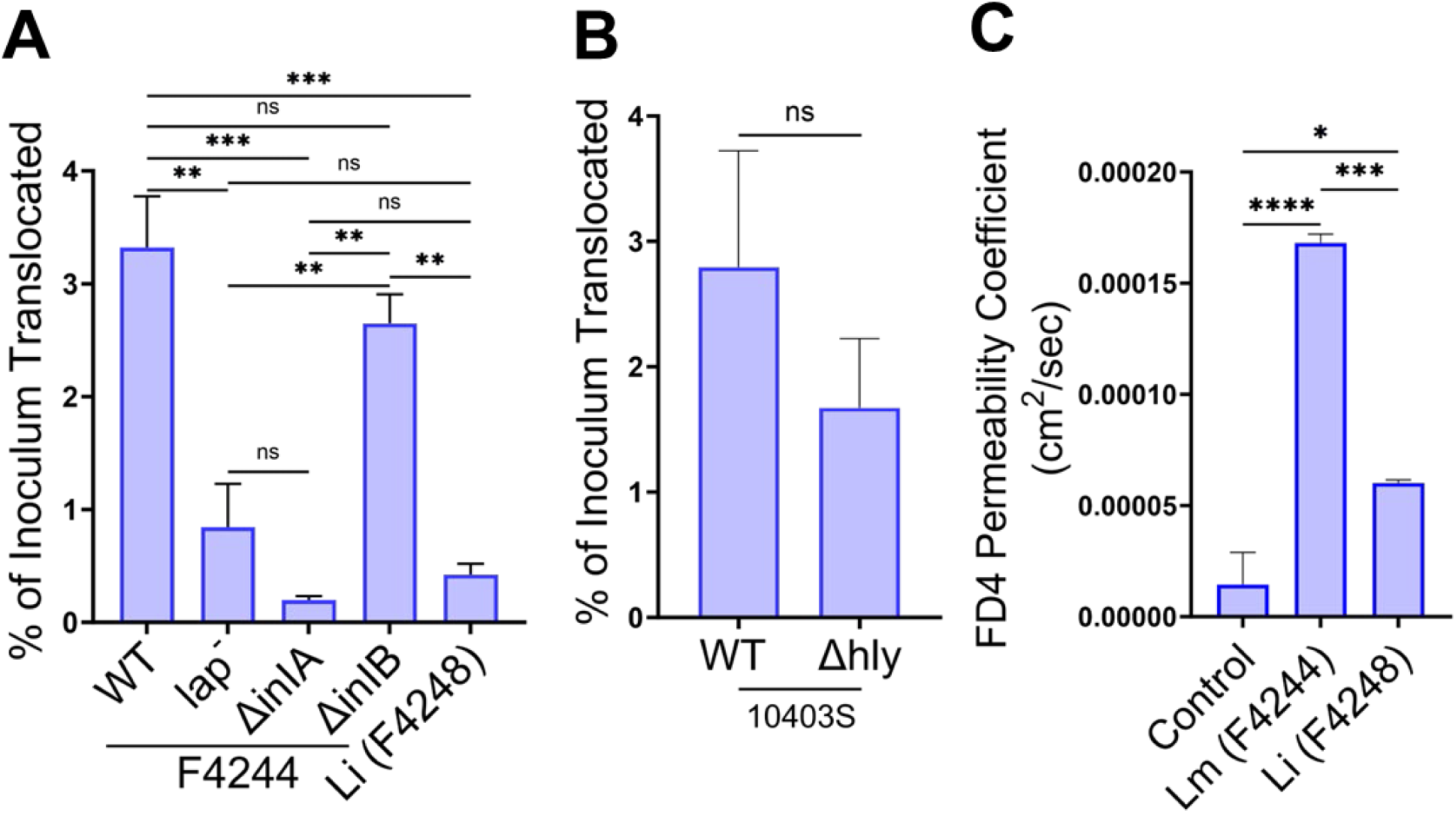
LAP and InlA facilitate extracellular translocation of *Listeria monocytogenes* across the placental co-culture model. Confluent BeWo–HPVECs co-cultures in Transwell inserts were apically inoculated with *L. monocytogenes* (*Lm*) or *L. innocua* (*Li*) at an MOI of 50 for 2 h at 37° C (A-C). (**A**) Translocated bacteria recovered from the basolateral chamber were quantified by plating and expressed as a percentage of the initial inoculum. Translocation was significantly reduced for *lap⁻*, *ΔinlA*, and *Li* compared with WT *Lm* F4244, whereas Δ*inlB* showed a modest, nonsignificant reduction. (**B**) Translocation comparison of WT *Lm* strain 10403S and its Δ*hly* mutant showed no significant difference in translocation efficiency. (**C**) Placental barrier permeability, assessed by FITC-dextran 4kDa (FD4) flux, was significantly increased following exposure to extracellular *Lm* compared with *Li*. Data in A–C are mean ± SEM from three independent experiments (n = 3 per treatment). Statistical significance was determined by one-way ANOVA with Tukey’s multiple-comparisons test. *, P < 0.05; **, P < 0.01; ***, P < 0.001. ****, P < 0.0001; ns, not significant.

Together, these findings demonstrate that LAP and InlA are essential mediators of efficient extracellular traversal of the placental barrier, whereas InlB and LLO contribute minimally in this context.

### Extracellular *Listeria monocytogenes* Enhances THP-1 Monocyte Transmigration

To model a physiological scenario in which *Lm* exists both extracellularly and within circulating monocytes, uninfected or *Lm*-infected THP-1 cells were co-incubated with extracellular *Lm* during transmigration. Extracellular *Lm* significantly increased transmigration of both uninfected THP-1 cells (3.97-fold) and previously infected THP-1 cells (2.69-fold) relative to unexposed controls, with uninfected cells exhibiting 47% greater migration than infected cells under these conditions (**Fig. 5A**). This condition also yielded the highest recovery of both extracellular and intracellular *Lm* from the basolateral chamber (**Fig. 5B**).

**FIG 5.**
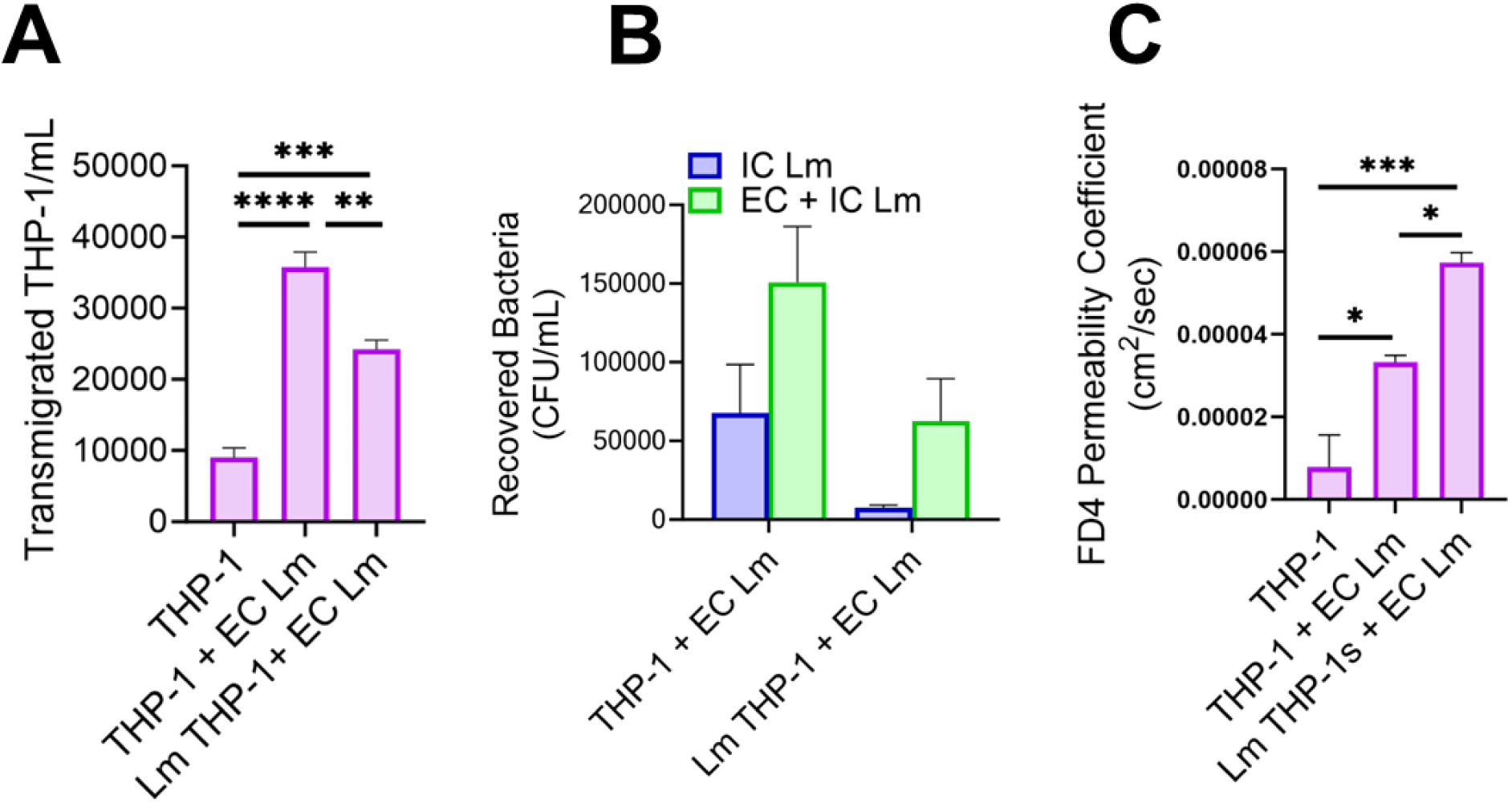
Effects of intracellular and extracellular *Lm* on THP-1 transmigration and placental barrier integrity in a co-culture model. Uninfected THP-1 monocytes (1 × 10⁶ cells) or THP-1 monocytes infected with *L. monocytogenes* (*Lm*; MOI 50, 1 h) were treated with gentamicin sulfate (50 µg/mL, 30 min), washed, and added to the apical chamber of BeWo–HPVECs co-culture Transwell inserts in F-12 medium. For conditions with extracellular *Lm* (EC Lm), co-cultures were apically inoculated with *Lm* (MOI 50) without gentamicin in either chamber. After 2 h at 37°C, basolateral compartments were collected to quantify transmigrated THP-1 cells, bacterial burdens, and barrier permeability. (**A**) Viable THP-1 monocytes recovered from the basolateral chamber, quantified by trypan blue exclusion and hemocytometer counting. (**B**) Bacterial burden recovered from the basolateral chamber, expressed as CFU/mL. Blue bars represent intracellular *Lm* recovered from transmigrated THP-1 cells, whereas green bars represent total *Lm* (intracellular plus extracellular bacteria). (**C**) Placental barrier permeability quantified as the FITC–dextran (4 kDa; FD4) permeability coefficient (cm²/s). Data in A–C represent mean ± SEM from three independent experiments (n = 3 per treatment). Statistical significance in A & C was determined by one-way ANOVA with Tukey’s multiple-comparisons test. *, P < 0.05; **, P < 0.01; ***, P < 0.001; ****, P < 0.0001.

Barrier permeability increased 4.24-fold with uninfected THP-1 cells exposed to extracellular *Lm* and 7.34-fold with infected THP-1 cells under the same conditions (**Fig. 5C**). Notably, despite showing lower transmigration, *Lm*-infected THP-1 cells induced substantially greater barrier permeability, indicating that infected monocytes exert more severe junction-disrupting effects per migrating cell.

Together, these data demonstrate that extracellular *Lm* enhances monocyte trafficking and markedly amplifies barrier disruption, supporting a synergistic contribution of extracellular and monocyte-associated bacteria to placental barrier compromise.

### LAP and InlB Drive Monocyte-Mediated Transmigration of *Lm* Across the Placental Barrier

To define how *Lm* virulence factors contribute to monocyte–mediated placental crossing, we first assessed uptake of WT strains (F4244, 10403S) and their Δ*inlA*, Δ*inlB*, *lap*⁻, and Δ*hly* derivatives in THP-1 monocytes. After 1 h of infection and 30 min of gentamicin treatment, only Δ*inlA* showed impaired internalization, whereas Δ*inlB*, *lap*⁻, and Δ*hly* maintained WT-like intracellular burdens (**Fig. 6A**). A short time-course assay (0.5 and 2.5 h gentamicin treatment) confirmed that all strains, aside from the expected Δ*inlA* uptake defect, maintained WT-like intracellular CFUs, indicating no early survival or replication impairment within monocytes (**Fig. S3A**). These data established that subsequent transmigration differences are not attributable to differential uptake.

**FIG 6.**
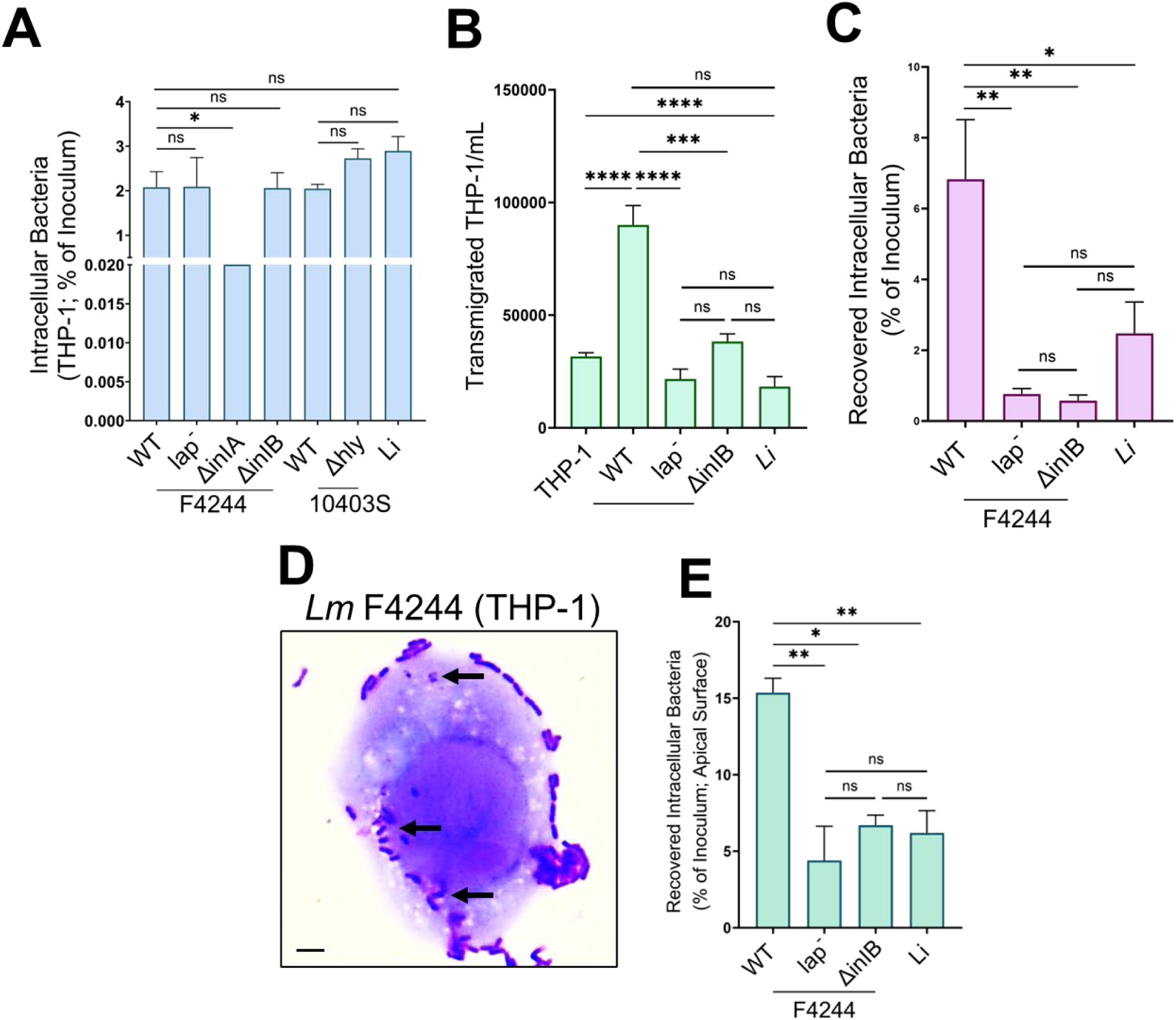
LAP- and InlB-dependent monocyte-mediated transmigration of *Listeria monocytogenes* across the placental co-culture model. (**A**) Intracellular uptake of *Listeria* strains by THP-1 monocytes. THP-1 cells (1 × 10⁶) were infected with WT *Listeria monocytogenes* (*Lm*) F4244, *lap⁻*, *ΔinlA*, *ΔinlB*, WT 10403S, *Δhly*, or *Listeria innocua* (*Li*) (MOI 50, 1 h) and treated with gentamicin sulfate (50 µg/mL, 30 min). Intracellular bacteria were quantified post-lysis and expressed as the percentage of the initial inoculum recovered. *ΔinlA* showed significantly reduced uptake. (**B**) Quantification of transmigrated THP-1 monocytes recovered from the basolateral chamber of the BeWo–HPVECs co-culture placental model after 2 h transmigration. THP-1 transmigration was significantly reduced for *lap⁻*, *ΔinlB*, and *Li* compared with WT *Lm* F4244. (**C**) Quantification of intracellular bacteria recovered from transmigrated THP-1 monocytes collected from the basolateral chamber after 2 h transmigration. Bacterial burdens were significantly reduced for *lap⁻*, *ΔinlB*, and *Li* relative to WT *Lm* F4244. (**D**) Representative Giemsa-stained Cytospins of gentamicin-protected THP-1 monocytes infected with WT *Lm* and recovered from the basolateral chamber after 2 h transmigration. Arrows indicate intracellular bacteria. Scale bar, 1 µm. (**E**) Invasion of apically attached monocytes and placental cells following transmigration. BeWo–HPVECs Transwells used for the 2 h transmigration assay were incubated for an additional 2 h in gentamicin-containing medium. Apical cells were collected, lysed, and intracellular bacteria enumerated. Invasion was significantly reduced for *lap⁻*, *ΔinlB*, and *Li* compared with WT *Lm* F4244. Data in A–C and E represent mean ± SEM from three independent experiments (n = 3 per treatment). Statistical significance was determined by one-way ANOVA with Tukey’s multiple-comparisons test. *, P < 0.05; **, P < 0.01; ***, P < 0.001; ****, P < 0.0001; ns, not significant. Exception: For WT (F4244) and *lap⁻* treatments in panel A, data represent five independent biological replicates (*n* = 5).

In the co-culture placental Transwell models, infections with *lap⁻* and Δ*inlB* caused 76% and 57% reductions in monocyte transmigration, respectively, relative to WT F4244 and was comparable to monocytes infected with *Li* (**Fig. 6B**). This impairment was accompanied by 89-92% decreases in intracellular bacteria recovered from the basolateral compartment (**Fig. 6C**). Giemsa-stained Cytospin preparations of the basolateral samples confirmed the presence of gentamicin-protected intracellular *Lm* within the transmigrated THP-1 cells (**Fig. 6D**). To quantify bacterial intracellular burdens in the apical adherent monocytes and the trophoblast layer, we applied an additional gentamicin step before lysis. WT F4244 yielded the highest intracellular CFUs, whereas *lap⁻* and *ΔinlB* showed 71% and 56% reductions, respectively, approaching levels observed in *Li* (**Fig. 6E**).

Deletion of *hly* in the 10403S background produced the opposite phenotype: Δ*hly* infection increased monocyte transmigration 2.9-fold and yielded fewer intracellular bacteria basolaterally (**Fig. S3B and S3C**), consistent with a model in which reduced LLO-mediated cytotoxicity may enhance monocyte survival and impaired phagosomal escape (21). Intracellular CFUs in the apical layers, comprising adherent monocytes and placental cells, were comparable between WT and Δ*hly* (**Fig. S3D**), indicating that LLO does not regulate apical entry into the barrier.

Together, these data demonstrate that LAP and InlB are dispensable for monocyte entry but function as key virulence determinants, enabling infected monocytes to traverse the placental barrier and deliver intracellular bacteria across it.

### *Lm*-Infected Monocytes Promote Pro-Transmigration Endothelial Phenotype

We next examined how *Lm*–infected monocytes influence placental endothelial activation. An adhesion-molecule array targeting 17 junctional and adhesion proteins was performed on harvested supernatants of placental co-cultures containing THP-1 monocytes under four conditions: untreated control, uninfected THP-1, *Lm*-infected THP-1, and *Li*-infected THP-1 (**Fig. 7A and 7B**). *Lm*-infected monocytes selectively upregulated BCAM (2.38-fold), P-selectin (2.25-fold) mediators of early rolling and adhesion, as well as VCAM-1 (1.73-fold), which supports firm attachment and transendothelial migration (**Fig. 7B and 7C**). Both *Lm* and *Li* triggered robust induction of ICAM-3, PECAM-1, NCAM-1, and NrCAM; however, consistently higher fold changes in *Lm*-infected samples, particularly for NCAM-1 and NrCAM, indicate stronger endothelial activation and barrier remodeling responses (**Fig. 7C, Table S1**). Collectively, adhesion-molecule profiling revealed a coordinated endothelial response aligned with sequential stages of monocyte recruitment: rolling, activation, firm adhesion, transmigration, and barrier modulation (**Table S1**).

**FIG 7.**
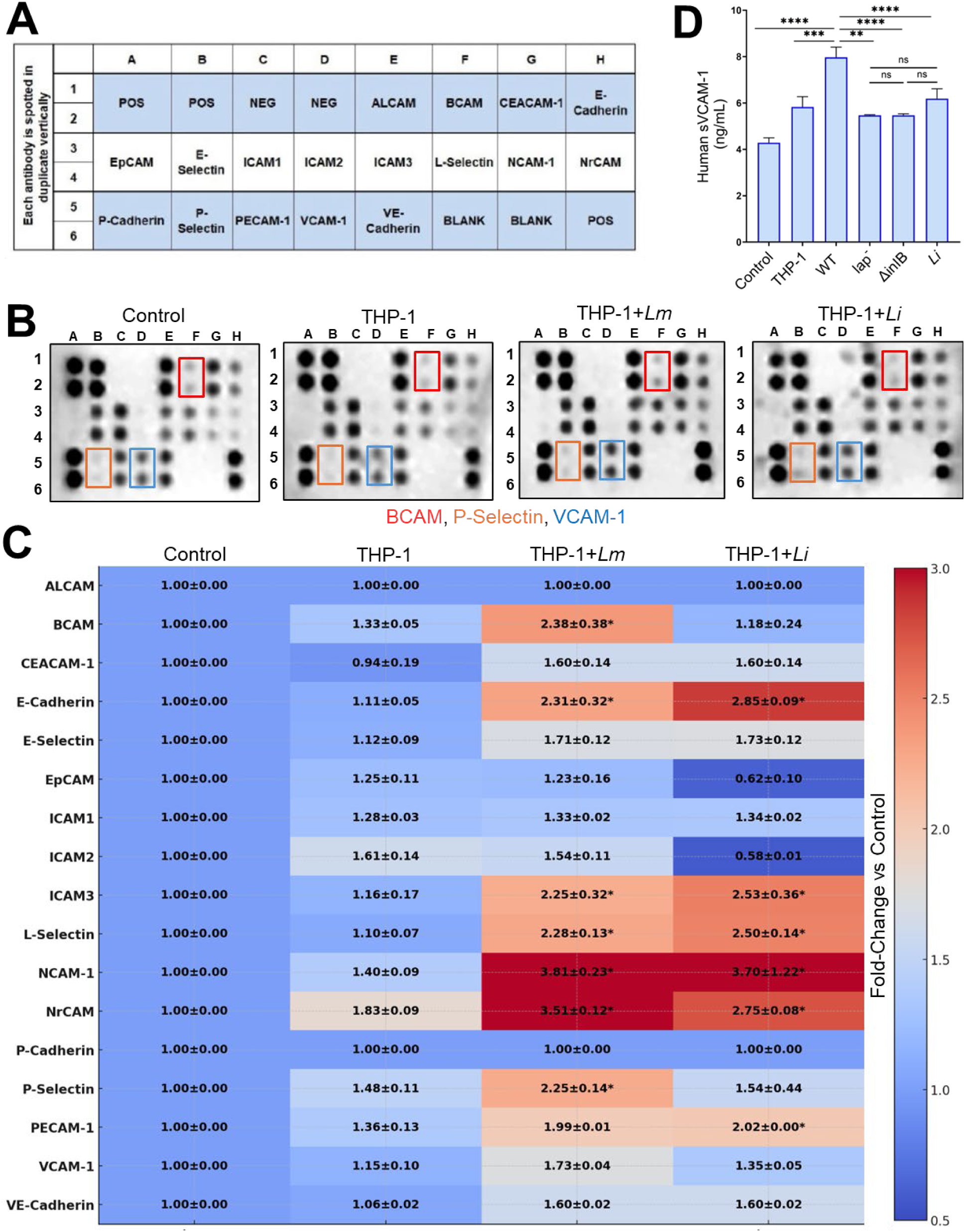
*Listeria monocytogenes* programs THP-1 monocytes toward a pro-transmigration endothelial phenotype. (**A**) Adhesion molecule array schematic. Apical supernatants were collected from BeWo–HPVECs placental co-culture barriers exposed to control conditions (no monocytes), uninfected THP-1 monocytes, *Listeria monocytogenes* (*Lm*)–infected THP-1 monocytes, or *Listeria innocua* (*Li*)–infected THP-1 monocytes and incubated on arrays containing 20 adhesion molecule–specific antibodies. (**B**) Representative adhesion-molecule array blots. Exposure to *Lm*-infected THP-1 monocytes resulted in marked upregulation of BCAM, P-selectin, and VCAM-1, whereas *Li*-infected THP-1 monocytes induced weaker or more selective changes. (**C**) Heatmap depicting fold-change expression of adhesion molecules relative to untreated no-monocyte control co-cultures. *Lm*-infected THP-1 monocytes selectively induced BCAM (∼2.4-fold), P-selectin (∼2.2-fold), and VCAM-1 (∼1.7-fold) compared with no-monocyte controls. Fold changes were calculated relative to untreated control co-cultures, normalized to internal positive controls. Data represent mean ± SD from pooled supernatants derived from four independent experiments per treatment, each analyzed in technical duplicate. Proteins exhibiting ≥2-fold induction relative to control are denoted with asterisks. (**D**) ELISA quantification of sVCAM-1 in apical supernatants. *Lm*-infected THP-1 monocytes induced a 1.86-fold increase in sVCAM-1 compared with control conditions. sVCAM-1 induction was significantly reduced with *lap⁻*, *ΔinlB*. Data represent mean ± SEM for atleast four independent samples (n = 4-5 per treatment). Statistical significance was determined by one-way ANOVA with Tukey’s multiple-comparisons test. ns, not significant; **, P < 0.01; ***, P < 0.001; ****, P < 0.0001.

Consistent with the array data, ELISA quantification confirmed a 1.86-fold *Lm-*specific induction of soluble VCAM-1 (sVCAM-1). This response was dependent on LAP and InlB, as *lap⁻* and Δ*inlB* mutants failed to upregulate sVCAM-1 (**Fig. 7D**). In contrast, soluble ICAM-1 upregulation was monocyte-dependent but independent of *Lm* infection (**Fig. S4**).

Together, these findings demonstrate that *Lm* reprograms the placental endothelium to promote monocyte adhesion and transmigration through selective LAP- and InlB-dependent induction of sVCAM-1, consistent with a mechanism that facilitates monocyte-mediated traversal of the placental barrier.

## DISCUSSION

The placenta is a formidable immunological and physical barrier, yet *Lm* can traverse it and cause severe fetal infection (7). Although InlA–E-cadherin interactions have been implicated in transplacental disease, whether *Lm* exploits monocytes to cross the placental barrier and whether internalin-independent virulence factors contribute to placental barrier crossing have remained unaddressed (4, 6, 8, 14, 32). Here, we validate a co-culture human placental Transwell model and define the virulence requirements for monocyte-assisted and extracellular traversal across the placental barrier. Our data support that LAP and InlB are indispensable for monocyte-mediated translocation, whereas LAP and InlA are required for extracellular traversal, revealing coordinated mechanisms that enable *Lm* to overcome trophoblast and endothelial defenses. Together, these results refine the framework of *Lm* placental translocation and identify LAP as a previously unrecognized virulence determinant central to both internalin-dependent and internalin-independent barrier crossing.

Our data suggest that the BeWo–HPVECs co–culture Transwell model recapitulates key features of the human placental barrier. Compared to single-cell monolayers, the co-culture exhibited reduced FITC-dextran permeability, higher TEER values, and well-defined tight junctions, reflecting enhanced barrier integrity. While *Lm* invasion into trophoblasts using gentamicin protection assay has been studied in Jeg-3 and BeWo cells, pathogens must cross both the trophoblast layer and the underlying fetal endothelium to reach the fetal circulation (14, 20, 33). This co-culture model captures these multistage translocation events, enabling dissection of pathogen-specific strategies for placental entry, overcoming limitations of single-layer *in vitro* systems, and providing a controlled, reproducible platform that complements *in vivo* models constrained by interspecies differences and ethical considerations. Although *in vivo* validation is ultimately required, our dual-cell system isolates trophoblast–endothelial interactions with a resolution that may not be achievable in animal models.

Our data reveal that *Li* undergoes unexpectedly similar uptake by THP-1 monocytes as *Lm*, despite lacking LIPI-1, likely reflecting passive phagocytosis. Unlike *Lm,* which represses flagellar expression at 37 °C, *Li* retains flagella, a feature that may promote recognition by host receptors or directly stimulate phagocytosis, thereby compensating for the absence of canonical invasion factors (34, 35). In contrast, consistent with previous observations, poor uptake of the *Lm ΔinlA* mutant suggests that InlA contributes to and compensates for monocyte interactions when flagella expression and flagellin-mediated adhesion are absent at host temperature (36). Notably, both *Lm* and *Li* displayed comparable early intracellular burdens, indicating that basic uptake and survival are not determinants of differential outcomes. Strikingly, however, only *Lm-*infected monocytes exhibited significantly enhanced transmigration across the placental barrier and increased delivery of viable intracellular bacteria, underscoring a virulence-dependent modulation of host cell trafficking. Together, these data highlight a critical distinction: while nonpathogenic *Li* can be phagocytosed and persist transiently, only pathogenic *Lm* actively reprograms monocyte adhesion and trafficking to promote placental barrier traversal. This supports a Trojan-horse mechanism defined here as the active transmigration of infected monocytes carrying viable intracellular bacteria.

Our study provides mechanistic insight into how *Lm* co-opts host monocytes to breach the placental barrier (**Fig. 8**). *Lm*-infected monocytes caused profound tight-junction disruption, marked by fragmentation of occludin and claudin-1, whereas uninfected monocytes produced only mild perturbations. Mechanistically, we show that LAP and InlB are indispensable for monocyte-assisted placental barrier translocation, with *lap* and *inlB* mutants exhibiting markedly reduced THP-1 transmigration and intracellular bacterial delivery despite normal uptake and early intracellular replication. These observations implicate LAP and InlB in post-entry events that govern monocyte adhesion, trafficking, and paracellular barrier crossing. Notably, enhanced monocyte adhesion and trafficking mediated by LAP and InlB may also potentiate an additional, previously proposed mechanism of placental infection—direct cell-to-cell spread from infected maternal phagocytes to trophoblasts (32). Once internalized by maternal monocytes, *Lm* escapes into the cytosol and uses ActA-driven actin-based motility to spread across membrane contacts, enabling transfer into trophoblasts without extracellular exposure (**Fig. 8)**.

**FIG 8.**
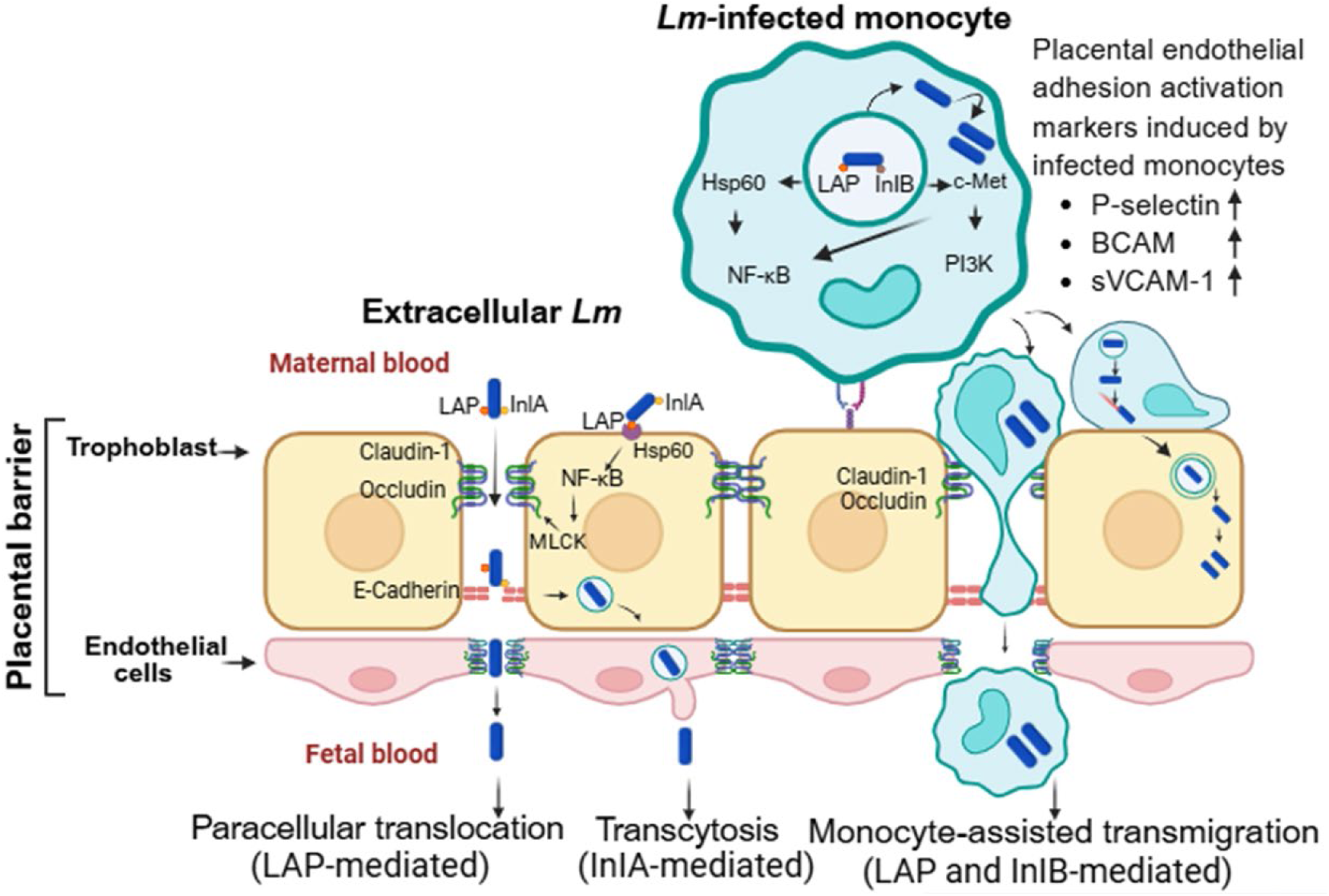
Proposed model of LAP- and internalin-dependent strategies for *Listeria monocytogenes* traversal of the human placental barrier. *Lm* crosses the placental barrier through two complementary routes: (left) extracellular translocation mediated by LAP and Internalin A (InlA), and (right) monocyte-mediated transmigration dependent on LAP and Internalin B (InlB). In the extracellular pathway, LAP–Hsp60 engagement at the trophoblast promotes paracellular barrier remodeling and translocation, with InlA-E-cadherin interaction additionally facilitating transcytosis across trophoblasts. In the monocyte-assisted pathway, intracellular *Lm* exploits circulating monocytes as Trojan-horse carriers. LAP–Hsp60 and InlB–c-Met signaling are proposed to activate NF-κB and PI3K/Akt pathways, thereby reprogramming infected monocytes toward a pro-transmigration phenotype. This monocyte-intrinsic activation is associated with the induction of placental endothelial adhesion and activation markers, including increased sVCAM-1, BCAM, and P-selectin. Enhanced endothelial activation promotes firm monocyte adhesion and paracellular migration and is accompanied by disruption of claudin-1 and occludin tight junctions. Increased monocyte retention and adhesion at the placental barrier may further facilitate ActA-mediated cell-to-cell spread, thereby amplifying bacterial dissemination across the barrier. Loss of LAP or InlA markedly impairs extracellular placental traversal, whereas loss of LAP or InlB selectively disrupts monocyte-mediated transmigration.

An intriguing question emerging from these findings is why *Lm* requires both LAP and InlB for monocyte-assisted bacterial delivery: do these factors provide partially redundant pro-adhesive cues, or does their cooperation create a signaling threshold that neither factor alone can achieve? LAP is an acetaldehyde alcohol dehydrogenase (AdhE). This moonlighting protein functions not only as a surface adhesin but also as a signaling scaffold within infected monocytes. LAP anchors to the *Lm* cell surface via electrostatic interactions with the glycine (G)–tryptophan (W) domain of InlB, thereby stabilizing its presentation and potentiating cell-intrinsic LAP–InlB signaling (1, 15). Although the LAP receptor Hsp60 is classically mitochondrial, we and others have demonstrated that *Lm* infection and inflammatory activation redistribute Hsp60 to the cell surface and cytosol, enabling autonomous signaling within infected monocytes (16, 17, 37–39). In parallel, c-Met is abundantly expressed on the surface of activated monocytes, providing a second cell-intrinsic signaling axis capable of engaging *Lm* InlB independently of endothelial contact (24). Consistent with this cooperative model, we previously showed that LAP and InlB, but not InlA, activate NF-κB signaling in RAW macrophages, supporting their broader role in modulating immune and barrier responses (17). We therefore propose that simultaneous engagement of Hsp60 and c-Met by the LAP–InlB complex creates a high-avidity signaling hub that activates NF-κB and PI3K/Akt pathways, priming infected monocytes to induce placental endothelial activation, including sVCAM-1 upregulation, thereby promoting firm adhesion and transmigration (**Fig.8**). In contrast, LLO was dispensable, underscoring the distinction between intracellular survival and barrier traversal. Together, these findings position LAP, scaffolded by InlB, as a noncanonical virulence module that reprograms monocyte adhesion and signaling pathways to enable monocyte-mediated crossing of the placental barrier.

Our data suggest that extracellular translocation of *Lm* across the placental barrier is critically dependent on LAP and InlA, with loss of either factor reducing traversal by 74–94% to *Li*–like levels, while InlB and LLO are dispensable. The minimal impact of *inlB* and *hly* deletion on placental traversal is consistent with prior observations that InlB- and LLO-deficient strains show negligible defects in BeWo trophoblast invasion, whereas InlA-deficient strains exhibit markedly reduced invasion, thereby confirming the established role of InlA and revealing LAP as an additional, previously unrecognized virulence determinant required for efficient extracellular barrier crossing (20). This phenotype aligns with *in vivo* studies establishing the essentiality of InlA–E-cadherin interactions for transplacental dissemination and raises the question of why *Lm* requires both LAP and InlA for efficient extracellular traversal (8, 14).

Our prior work at the intestinal barrier demonstrated that LAP-mediated remodeling of apical epithelial junctions is the initiating event in a two-step process that (i) opens paracellular routes via LAP–Hsp60 signaling and (ii) exposes normally shielded basolateral E-cadherin to enable InlA-dependent epithelial transcytosis, while LAP–Hsp60 can also drive internalin-independent paracellular translocation (17, 18). We propose that an analogous cascade operates at the placenta: trophoblast layers express high levels of Hsp60, and LAP binding to trophoblast Hsp60 may remodel junctional complexes and promote paracellular leakage, thereby unmasking basolateral E-cadherin and providing InlA access to its receptor for efficient transcytosis across the trophoblast layer (40, 41) (**Fig. 8**). This model explains the cooperative requirement for LAP and InlA, integrates receptor engagement at both Hsp60 and E-cadherin, and is further supported by experimental and clinical evidence that severe neonatal listeriosis can occur even with internalin A-deficient strains, underscoring the LAP-mediated, internalin-independent paracellular translocation route that broadens the arsenal of *Lm* strategies for breaching host barriers (**Fig. 8**). (32, 42, 43).

Although this co-culture placental model captures essential trophoblast–endothelial interactions with a level of mechanistic resolution that may not be achievable in animal systems, it does not fully recapitulate the architectural or immunological complexity of the *in vivo* placenta. *In vivo* studies or primary maternal monocytes will ultimately be required to confirm the relative contributions of LAP, InlA, and InlB under physiological conditions. In addition, while our adhesion-molecule profiling implicates VCAM-1 and related pathways in promoting monocyte trafficking, future studies will be needed to determine the functional requirement of these receptors during placental transmigration.

In conclusion, our work uncovers two complementary strategies by which *Lm* breaches the human placental barrier: monocyte–mediated translocation driven by LAP and InlB and direct extracellular invasion requiring LAP and InlA (**Fig. 8**). By assigning route-specific functions to these virulence factors, we reveal LAP as a critical, non-canonical determinant of placental barrier traversal and a potential driver of vertical transmission and highlight the LAP–InlB axis as a potential therapeutic target. Targeting LAP-mediated adhesion or LAP–InlB signaling may therefore represent a route-selective strategy to block placental invasion. Beyond advancing our understanding of listerial pathogenesis in pregnancy, the co-culture placental model provides a versatile platform to dissect host–pathogen interactions and to accelerate the development of interventions against *Lm* and other vertically transmitted pathogens.

## MATERIALS AND METHODS

### Cell Culture

ATCC BeWo human placental epithelial cells (catalog no. CCL-98) were cultured in F-12K medium supplemented with 10% (v/v) fetal bovine serum (FBS). Primary human placental vascular endothelial cells (HPVECs) (ScienCell, catalog no. 7100) were cultured in F-12K medium supplemented with 10% (v/v) FBS and 1% endothelial cell growth supplement (ECGS; ScienCell, catalog no. 1052) and used between passages 3 and 7. ATCC THP-1 human monocytic leukemia cells (catalog no. TIB-202) were maintained in RPMI-1640 medium supplemented with 0.05 mM 2-mercaptoethanol and 10% (v/v) FBS. All cell lines were maintained in a humidified incubator at 37°C with 5% CO₂. Unless otherwise stated, cells were maintained without antibiotics. Cell line sources, catalog numbers, and culture reagents are listed in Table S2 (Key Resources).

### Bacterial Cultivation

*L. monocytogenes* wild-type F4244 (serotype 4b, clinical isolate, CC6, Lineage I) and respective isogenic mutants (Δ*inlA*, Δ*inlB*, and *lap−*) and a GFP-expressing F4244 strain maintained in 2µg/mL erythromycin; the wild type of *L. innocua* (F4248); *L. monocytogenes* wild-type 10403s (serotype 1/2a, skin isolate, CC7, Lineage II) and its respective Δ*hly* mutant strain were cultured in tryptic soy broth (TSB) containing 0.6% yeast extract (YE). All strains were cultured at 37°C with agitation (120 rpm on an orbital shaker) for 14–18 h except the *lap−* insertion mutant strain, which was grown in TSB-YE supplemented with 5 µg/mL erythromycin and cultured at 42°C with agitation (120 rpm) for 14–18 h. Detailed strain information is provided in Table S2 (Key Resources).

### Human Placental Barrier Co-Culture Model

A dual-layer human placental co-culture model was established using polyethylene terephthalate (PET) Transwell inserts (Corning). For barrier formation and permeability assays, Transwell inserts with 4.0-µm pore size were used. Transwell membranes were coated with bovine plasma fibronectin at a concentration of 15 µg/mL in Dulbecco’s phosphate-buffered saline (DPBS). The underside of the inserts was coated for at least 2 h at room temperature.

Primary HPVECs were seeded on the underside of the inserts at densities of 2.8 × 10⁴ cells per insert (12-well format) or 8.25 × 10³ cells per insert (24-well format). After allowing 1 h for attachment, inserts were inverted, and BeWo cells were seeded on the apical surface at identical densities. Co-cultures were maintained in F-12K medium supplemented with 10% FBS and 1% ECGS at 37°C with 5% CO₂ and allowed to mature for at least 72 h prior to experimentation.

### Transepithelial Electrical Resistance (TEER) Measurement

Transepithelial electrical resistance (TEER) was used to assess barrier integrity and tight junction robustness in the BeWo–HPVECs dual-layer placental co-culture mode l(17–19). TEER measurements were performed using a Millicell ERS-2 voltohmmeter (Millipore) with electrodes positioned on both the apical and basolateral compartments of the Transwell inserts. Resistance measurements were obtained for blank inserts (no cells), BeWo monocultures, HPVECs monocultures, and BeWo–HPVECs co-cultures. Raw resistance values were corrected by subtracting the resistance of blank inserts and normalized to membrane surface area. Final TEER values were expressed as Ω·cm².

### FITC-Dextran Permeability Assay

A 1 mg/mL fluorescein isothiocyanate–dextran (4 kDa; FD4) working solution was prepared in F-12K medium. FD4 was added to the apical compartment of Transwell inserts containing BeWo monocultures, HPVECs monocultures, or BeWo–HPVECs co-cultures and incubated at 37°C with 5% CO₂ (17–19). For barrier integrity and transmigration assays, FD4 was present throughout the 2 h incubation period. In transmigration experiments, the apical compartment additionally contained gentamicin sulfate (50 µg/mL) and either uninfected or *Lm*-infected THP-1 monocytes (1×10⁶ cells/well), as indicated, except under conditions where extracellular *Lm* was added.

At designated time points, 50 µL samples were collected from the basolateral compartment and transferred to a black 96-well plate. Samples were immediately replaced with fresh medium to maintain constant volume. Fluorescence was measured using a Tecan Genios microplate reader (excitation 485 nm, emission 528 nm). FD4 concentrations were calculated from a standard curve. Permeability coefficients (Pc) were calculated using the equation: *Pc* = (*Vr* × *Cf*)/(*Ci* × *A* × *t*) where *Vr* is the receiver volume (mL), *Cf* is the final FD4 concentration in the basolateral compartment (mg/mL), *Ci* is the initial FD4 concentration in the apical compartment (mg/mL), A is the membrane surface area (cm²), and t is time (seconds).

### Preparation of THP-1 Cells Harbouring Internalized *Listeria* spp

To generate THP-1 monocytes harbouring internalized bacteria, THP-1 cells were infected with the following *Listeria* strains at a multiplicity of infection (MOI) of ∼50: *Li* F4248; *Lm* F4244 and its isogenic mutants (*ΔinlA*, *ΔinlB*, and *lap⁻*); and *Lm* 10403S and its isogenic *Δhly* mutant. Infections were carried out for 1 h at 37°C.To eliminate extracellular bacteria, infected THP-1 cells were treated with gentamicin sulfate (50 µg/mL) for 0.5 or 2.5 h at 37°C, as indicated. Cells were then washed thoroughly with DPBS and lysed using 0.1% Triton X-100. Intracellular bacterial burdens were quantified by serial dilution of cell lysates and plating on tryptic soy agar supplemented with 0.6% yeast extract (TSA-YE) plates for CFU enumeration following incubation at 37°C for 48 h. Intracellular recovery was expressed as the percentage of the initial inoculum, calculated using the formula: Percent intracellular recovery= (CFUs recovered from the lysate / total CFUs in the inoculum added to THP-1 monocytes) × 100

### Quantification of Internalized Bacteria in THP-1 Monocytes by Giemsa Staining

Following infection and gentamicin sulfate (50 µg/mL) protection as described above, intact THP-1 monocytes harboring internalized *Lm* or *Li* were processed for cytospin preparation and Giemsa staining prior to cell lysis. Briefly, 200 µL of THP-1 cell suspension containing approximately 1 × 10⁵ cells was loaded into Cytofunnel assemblies and centrifuged using an Epredia Cytospin™ 4 cytocentrifuge. Cytospins were generated using Program 1 (1500 rpm for 8 min) to deposit cells evenly onto poly-L-lysine–coated microscope slides (Polyssine, white).

Immediately following centrifugation, slides were fixed in 4% formaldehyde prepared in 1× PBS for 15–20 min at room temperature, followed by three gentle washes with PBS (5 min per wash) to remove residual fixative. Slides were then stained with Giemsa solution for 2–3 min, rinsed twice with deionized water, and air-dried. Stained cytospins were examined using a bright-field microscope (ECHO; Model REB-01-E2) equipped with a 100× oil-immersion objective (Plan Achromat, numerical aperture 1.25, working distance 0.21 mm). Gentamicin-protected cell-associated bacteria were manually quantified across multiple randomly selected fields, yielding a total of 150–175 THP-1 cells analyzed per treatment condition. In select experiments, cytospins were also prepared from basolateral compartment samples collected following transmigration assays and processed identically to confirm that intracellular gentamicin protected *Lm* within transmigrated THP-1 monocytes.

### Assessment of Transmigration by *Listeria*-Infected THP-1 Monocytes

To assess the ability of *Lm* or *Li*–infected THP-1 monocytes to transmigrate across the dual-layer placental Transwell model, 8.0-µm pore size Transwell inserts (Corning) were used to permit active leukocyte migration. Briefly, 1 × 10⁶ uninfected or infected THP-1 cells were added to the apical compartment of each insert in 150 µL of culture medium. Unless otherwise indicated, F-12K medium containing gentamicin sulfate (50 µg/mL) was added to both the apical and basolateral compartments to eliminate extracellular bacteria during the transmigration period. Transwell plates were incubated for 2 h at 37°C in a humidified incubator with 5% CO₂.

Following incubation, media from both the apical and basolateral compartments were collected. Apical supernatants were stored at –80°C for subsequent analysis by ELISA. Basolateral contents were processed to quantify both transmigrated THP-1 monocytes and intracellular bacteria. The entire basolateral volume was centrifuged at 900 rpm for 5 min to pellet transmigrated cells and associated bacteria. Pellets were washed twice with DPBS and resuspended in 300 µL F-12K medium.

To quantify transmigrated THP-1 monocytes, 100 µL of the suspension was stained with Trypan Blue and viable cells were counted using a hemocytometer, based on cell size and morphology. The remaining 200 µL was used to quantify intracellular bacteria. THP-1 cells were lysed with 0.1% Triton X-100 for 15 min, and lysates were serially diluted and plated on TSA-YE plates. Plates were incubated at 37°C for 48 h, after which CFUs were enumerated.

Transmigration efficiency was quantified as the number of THP-1 monocytes recovered per milliliter of basolateral medium. Intracellular bacterial delivery across the placental barrier was quantified either as intracellular CFUs recovered from lysates of transmigrated THP-1 monocytes per milliliter of basolateral medium or as the percentage of the initial intracellular inoculum within THP-1 monocytes recovered in transmigrated cells. The latter was calculated as: (CFUs recovered from lysates of transmigrated THP-1 monocytes/total intracellular CFUs in the initial THP-1 inoculum) × 100.

For conditions involving extracellular *Lm* exposure, gentamicin sulfate was omitted from both apical and basolateral compartments, and extracellular infection was performed at a multiplicity of infection (MOI) of ∼50. Extracellular bacteria were quantified by collecting culture supernatants from the apical or basolateral compartments, as indicated for each experiment. Collected media were centrifuged at 900 rpm for 5 min to remove host cells and debris, and the clarified supernatants were serially diluted in sterile phosphate-buffered saline (PBS) and plated TSA-YE. Plates were incubated at 37 °C for 48 h, and CFUs were enumerated.

Intracellular bacteria were quantified following gentamicin treatment (50 µg/mL) to eliminate extracellular bacteria. After antibiotic exposure, cells were washed thoroughly three times with PBS to remove residual gentamicin. Host cells were then lysed using 0.1% Triton X-100 in PBS for 15 min at room temperature to release intracellular bacteria. Lysates were serially diluted and plated on TSA-YE plates, incubated at 37 °C for 48 h, and CFUs were counted. Total bacterial burden was calculated as the sum of intracellular and extracellular CFUs recovered from the corresponding lysates and supernatants for each condition.

### Assessment of Bacterial Invasion at the Apical Placental Barrier

Following completion of the transmigration assay, bacterial invasion of apically attached monocytes and placental cells was assessed using the same Transwell inserts. To remove nonadherent and loosely bound cells, the apical surface was gently washed twice with DPBS. F-12K medium containing gentamicin sulfate (50 µg/mL) was then added to the apical chamber (150 µL per insert), and the inserts were incubated for an additional 1 h at 37 °C with 5% CO₂. After incubation, the apical surface was washed twice with DPBS to remove residual gentamicin (17–19).

Cells adherent to the apical surface were lysed by addition of 150 µL of 0.1% Triton X-100 for 15 min to release intracellular bacteria. Lysates were collected, serially diluted, and plated on TSA-YE plates to enumerate viable intracellular CFUs, representing bacterial invasion of placental cells and attached monocytes. Intracellular recovery was expressed as the percentage of the initial intracellular inoculum within THP-1 monocytes, calculated using the following formula: (CFUs recovered from apical lysates / total CFUs of the intracellular inoculum within THP-1 monocytes) × 100.

### Fluorescence-Based Confirmation of Intracellular Bacteria During Transmigration

In select experiments, THP-1 monocytes were labelled using the Cytopainter Cell Tracking Staining Kit (red fluorescence) according to the manufacturer’s instructions. Labeled cells were infected with GFP-expressing *Lm* at an MOI of ∼50 for 1 h, followed by gentamicin treatment (50 µg/mL for 30 min) to eliminate extracellular bacteria. Cytopainted THP-1 cells were examined before and after transmigration by collecting samples from the basolateral compartment. Fluorescence imaging was performed using a Zeiss LSM 800 confocal laser-scanning microscope equipped with 405 nm, Argon, and 561 nm lasers. Images were acquired using a 63× oil-immersion objective (NA 1.40). Z-stack images were collected at 0.25 µm intervals, and X–Z and Y–Z orthogonal reconstructions were generated to confirm intracellular localization of GFP-*Lm* within transmigrated THP-1 monocytes.

### Immunostaining of Cell–Cell Junctions

Cells cultured on Transwell insert membranes were fixed with 4% formaldehyde in PBS for 20 min at room temperature. Inserts were washed three times with PBS (5 min per wash) and incubated in blocking buffer for 1 h at room temperature. All antibodies and reagents are listed in the Key Resources Table (Table S2). Mouse anti-occludin or anti–claudin-1 primary antibodies were diluted 1:100 in antibody dilution buffer (ABDB), and inserts were incubated with the primary antibody solution overnight (18 h) at 4°C (17–19). Following incubation, inserts were washed five times with PBS (5 min per wash) and subsequently incubated with goat anti-mouse secondary antibodies conjugated to Alexa Fluor 488 or Alexa Fluor 555, diluted 1:500 in ABDB, for 1 h at room temperature. Inserts were then washed three times with PBS (5 min per wash). Nuclei were counterstained with DAPI (1% solution prepared in PBS) for 10 min at room temperature in the dark. Inserts were washed once with PBS, membranes were carefully excised using a razor blade, and mounted onto glass microscope slides using ProLong Gold Antifade mounting medium. Coverslips were applied and allowed to cure for at least 1 h prior to imaging. Confocal imaging was performed using a Zeiss LSM 800 laser-scanning confocal microscope equipped with 405 nm, Argon, and 561 nm lasers. Images were acquired using a 63× oil-immersion objective (NA 1.40). Z-stack images were collected at 0.25 µm intervals across ∼10 µm–thick co-cultures, and X–Z and Y–Z orthogonal reconstructions were generated using Zeiss ZEN software.

### Extracellular Bacterial Translocation

*Lm* and *Li* strains were cultured overnight as described above. Bacterial density was estimated by measuring optical density at 600 nm (OD₆₀₀) and confirmed by plating. Bacteria were resuspended in serum-free DMEM and added directly to the apical compartment of Transwell inserts containing the BeWo–HPVECs dual-layer placental co-culture model at a multiplicity of infection (MOI) of ∼50 (17–19). Following 2 h incubation at 37°C in a humidified incubator with 5% CO₂, media from the basolateral compartment were collected and subjected to serial dilution plating on TSA-YE plates. Plates were incubated at 37°C for 48 h, after which colony-forming units (CFUs) were enumerated. Translocation efficiency was calculated as the percentage of the initial inoculum recovered in the basolateral compartment.

### Human Cell Adhesion Molecule Array

Secreted adhesion molecules were profiled using the RayBio® C-Series Human Cell Adhesion Molecule Array 1 (Cat. No. AAH-CAM-1; RayBiotech) according to the manufacturer’s instructions. Apical conditioned supernatants were collected from BeWo–HPVECs placental co-culture barrier models under four experimental conditions: untreated control (no monocytes), uninfected THP-1 monocytes, THP-1 monocytes infected with *Lm*, or THP-1 monocytes infected with *Li*. Supernatants from four independent experiments per condition were pooled and clarified by centrifugation to remove cellular debris.

Briefly, cleared supernatants were incubated overnight at 4 °C with pre-blocked array membranes containing 20 pre-immobilized human adhesion molecule–specific capture antibodies. Membranes were subsequently incubated with a biotinylated detection antibody cocktail followed by horseradish peroxidase (HRP)–conjugated streptavidin. Chemiluminescent signals were developed using substrate reagents provided with the kit. Membranes were imaged using a Bio-Rad chemiluminescence imaging system and analyzed with Image Lab software (Bio-Rad Laboratories). Spot intensities were quantified using ImageJ software (NIH), background-subtracted, and normalized to internal positive controls on each membrane. Fold-change values were calculated relative to untreated control co-cultures.

### Human sVCAM-1 and sICAM-1 ELISA

Levels of soluble vascular cell adhesion molecule-1 (sVCAM-1) in apical supernatants were quantified using a Human sVCAM-1 ELISA kit (Invitrogen; Cat. no. KHT0601) according to the manufacturer’s instructions. Briefly, sVCAM-1 standards were reconstituted to a starting concentration of 75 ng/mL and serially diluted to generate a standard curve. Apical conditioned supernatants (100 µL per well) were added to 96-well plates pre-coated with anti-human sVCAM-1 capture antibody. Following the addition of the biotin-conjugated detection antibody, plates were incubated for 2 h at 37 °C and washed according to kit instructions. Streptavidin-HRP was then added and incubated for 30 min at room temperature. After washing, chromogenic substrate was added, and absorbance was measured at 450 nm using a microplate reader. sVCAM-1 concentrations in experimental samples were calculated from the standard curve.

Soluble intercellular adhesion molecule-1 (sICAM-1) levels were measured using a RayBio® Human sICAM-1 ELISA kit (RayBiotech; Cat no. ELH-ICAM1-1) following the manufacturer’s protocol. Apical supernatants were diluted in Assay Diluent B as recommended. Standards were prepared by serial dilution of the supplied positive control. Experimental samples (100 µL per well) were added to antibody-coated 96-well plates and processed according to kit instructions. Absorbance was measured at 450 nm, and sICAM-1 concentrations were calculated from the standard curve.

### Statistical Analysis

All data were analyzed using GraphPad Prism software. Results are presented as the mean ± standard error of the mean (SEM) unless otherwise noted from at least three independent biological experiments, with sample sizes (n) indicated in the corresponding figure legends. Comparisons between two groups were performed using an unpaired two-tailed Student’s t-test. Comparisons among multiple groups were conducted using one-way or two-way analysis of variance (ANOVA), followed by Tukey’s multiple-comparisons post hoc test.

## DATA AVAILABILITY

All strains used in this study will be made available upon publication and upon request.

## ACKNOWLEDGMENTS

The research in our laboratory was supported by start-up funds at Old Dominion University (ODU), as well as funding from the USDA National Institute of Food and Agriculture (NIFA) (Award No. 2023-67017-40051 to R.D.) and the Commonwealth Health Research Board (CHRB) (Award No. 221-01-25 to R.D.). A.A. and R.D. gratefully acknowledge Drs. Wayne Hynes, Lisa Schollenberger, and Isaura Simões for their critical input, insightful discussions, and valuable feedback during the development of this work.

## AUTHOR CONTRIBUTIONS

A.A. and B.A. contributed equally to this work and designed the experiments, performed the experiments, interpreted the data, and wrote parts of the manuscript. R.D. generated the idea, wrote the first draft of the manuscript, designed and performed the experiments, interpreted the data, revised the manuscript, and acquired funding. All other authors (Z.A.H., D.B., O.R.O., and M.P.) assisted in conducting experiments. All authors reviewed the manuscript.

## SUPPLEMENTAL FIGURES AND TABLES

**Figure S1.**
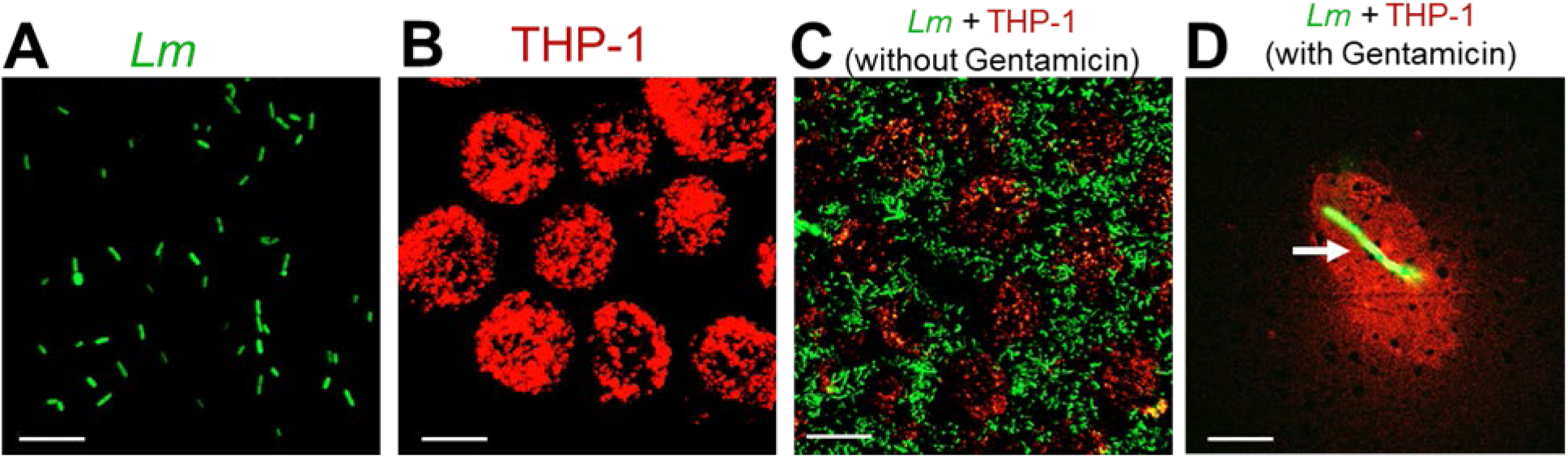
Representative Confocal fluorescence microscopy of GFP-expressing *Listeria monocytogenes* (*Lm*) within CellTracker Red–labeled THP-1 monocytes. (**A**) GFP-expressing *Lm*. Scale bar, 5 µm. (**B**) THP-1 monocytes labeled with CellTracker™ Red dye. Scale bar, 5 µm. (**C**) THP-1 monocytes infected with GFP-expressing *Lm* (MOI 50, 1 h) prior to gentamicin treatment. (**D**) Representative cell showing intracellular *Lm* following gentamicin protection (50 µg/mL, 30 min). Arrows indicate gentamicin-protected, cell-associated/intracellular *Lm*. Scale bar, 5 µm.

**Figure S2.**
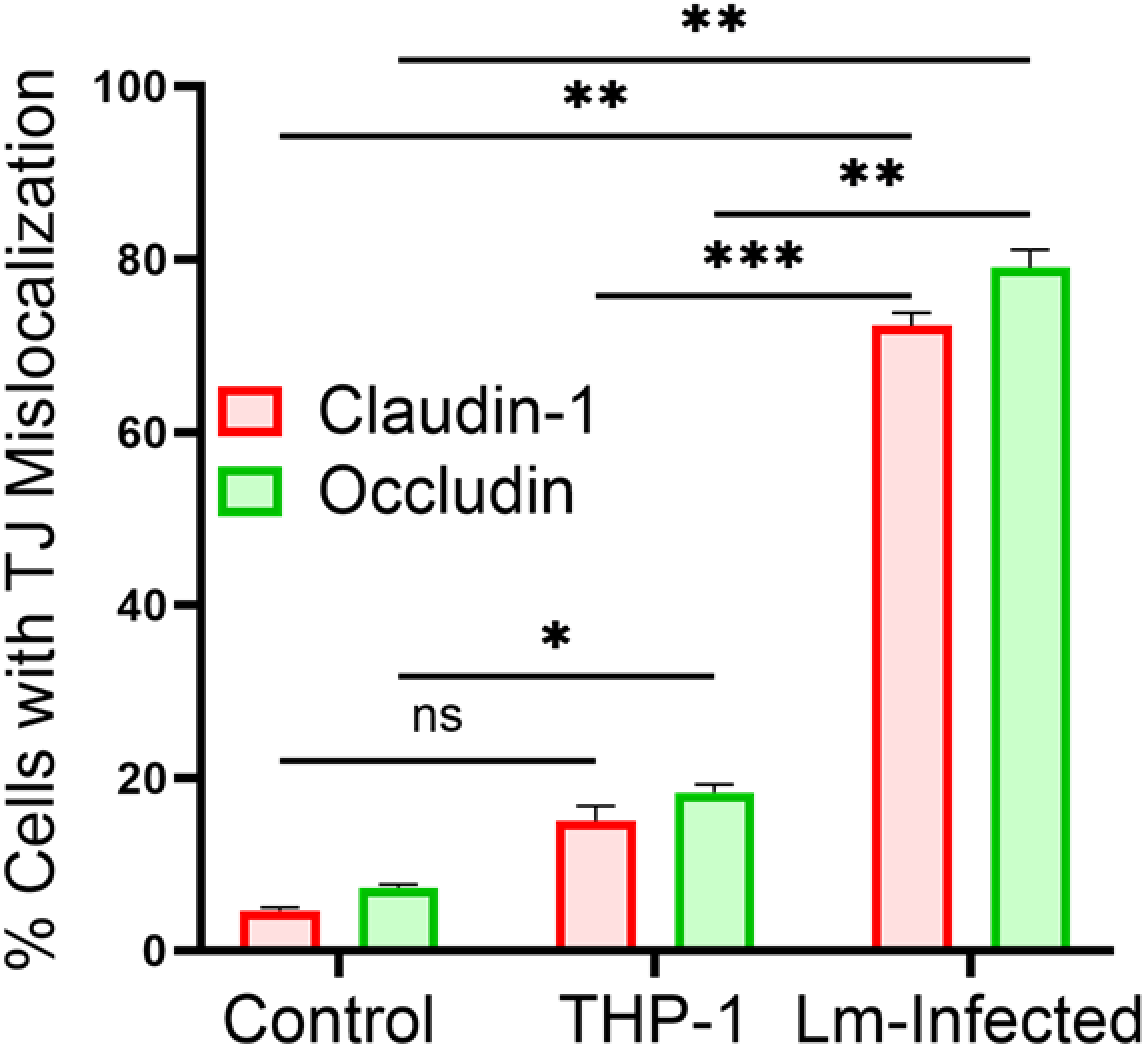
*Lm*–infected monocytes induce mislocalization of placental barrier tight junction proteins. Quantitative analysis of claudin-1 and occludin junctional mislocalization in BeWo–HPVECs co-cultures under three conditions: untreated control, exposure to uninfected THP-1 monocytes, or exposure to *Lm*–infected THP-1 monocytes. Each biological replicate consisted of five randomly selected fields per condition, with 8–10 cells analyzed per field. Data are presented as mean ± SEM from three independent experiments (n = 3). Statistical significance was determined by two-way ANOVA with Tukey’s multiple-comparisons test. ns, not significant; *, P < 0.05; **, P < 0.01; ***, P < 0.001.

**Figure S3.**
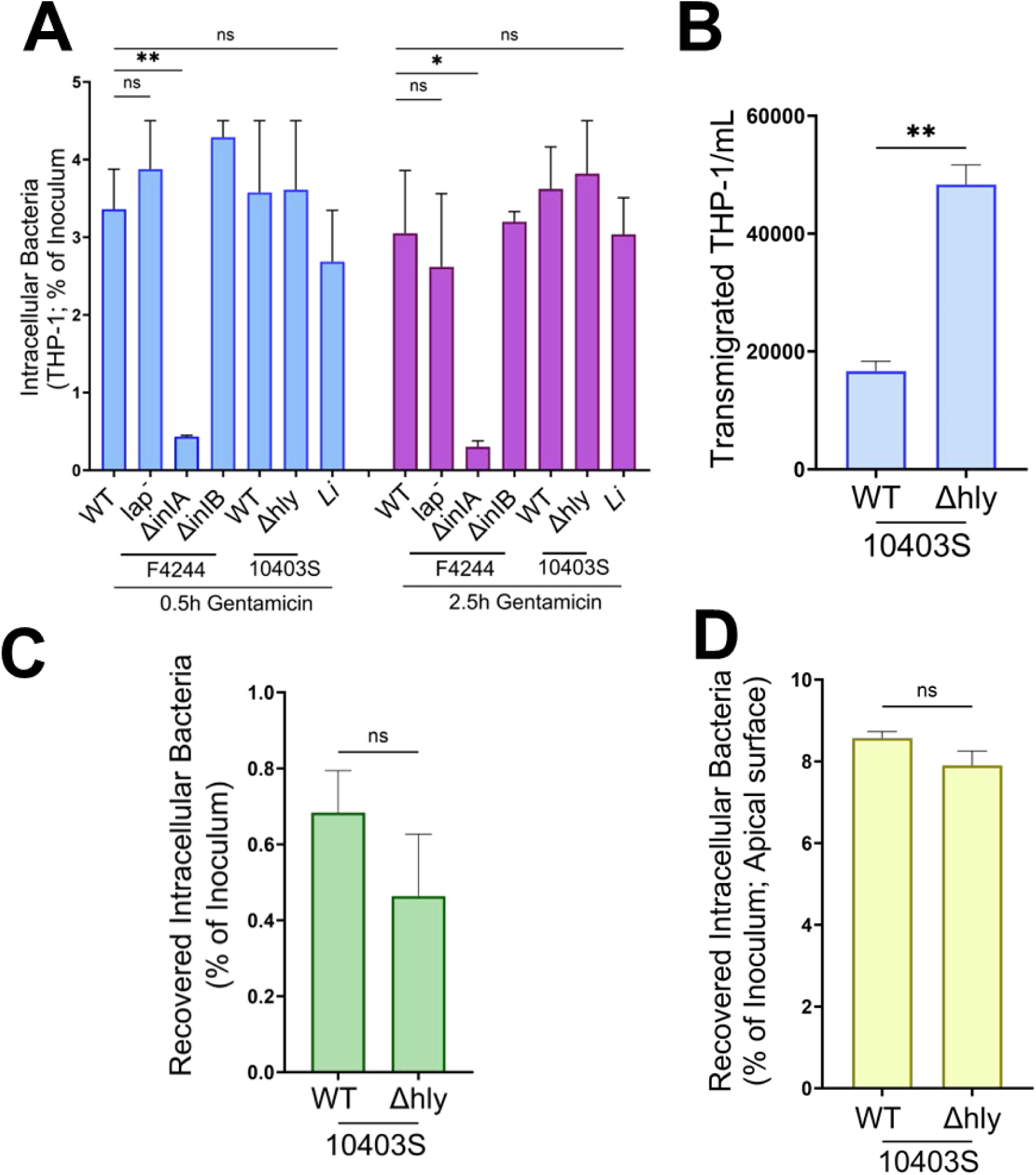
Intracellular uptake and effect of Listeriolysin O (LLO) on monocyte-mediated transmigration of *Lm*. (**A**) Intracellular uptake of *Listeria* strains by THP-1 monocytes. THP-1 cells (1 × 10⁶) were infected with WT *Lm* F4244, *lap⁻*, *ΔinlA*, *ΔinlB*, WT 10403S, *Δhly*, or *Listeria innocua* (*Li*) (MOI 50, 1 h) and treated with gentamicin sulfate (50 µg/mL) for 30 min or 2.5h. Intracellular bacteria were quantified post-lysis and expressed as the percentage of the initial inoculum recovered. *ΔinlA* exhibited significantly reduced uptake relative to WT strains. (**B**) Quantification of transmigrated THP-1 monocytes recovered from the basolateral chamber of the BeWo–HPVECs co-culture placental model after 2 h of transmigration. THP-1 transmigration was significantly increased for *Δhly* relative to WT *Lm* 10403S. (**C**) Quantification of intracellular bacteria recovered from transmigrated THP-1 monocytes collected from the basolateral chamber after 2 h transmigration. (**D**) Invasion of apically attached monocytes and placental cells following transmigration. BeWo–HPVECs Transwells used for the 2 h transmigration assay were incubated for an additional 2 h in gentamicin-containing medium. Apical cells were collected, lysed, and intracellular bacteria enumerated. Data in **A**–**D** represent mean ± SEM from three independent experiments (n = 3 per treatment). Statistical significance for panel A was determined using one-way ANOVA with Tukey’s multiple-comparisons test. Panels B–D were analyzed using an unpaired two-tailed Student’s t test. ns, not significant; *, P < 0.05; **, P < 0.01.

**Figure S4:**
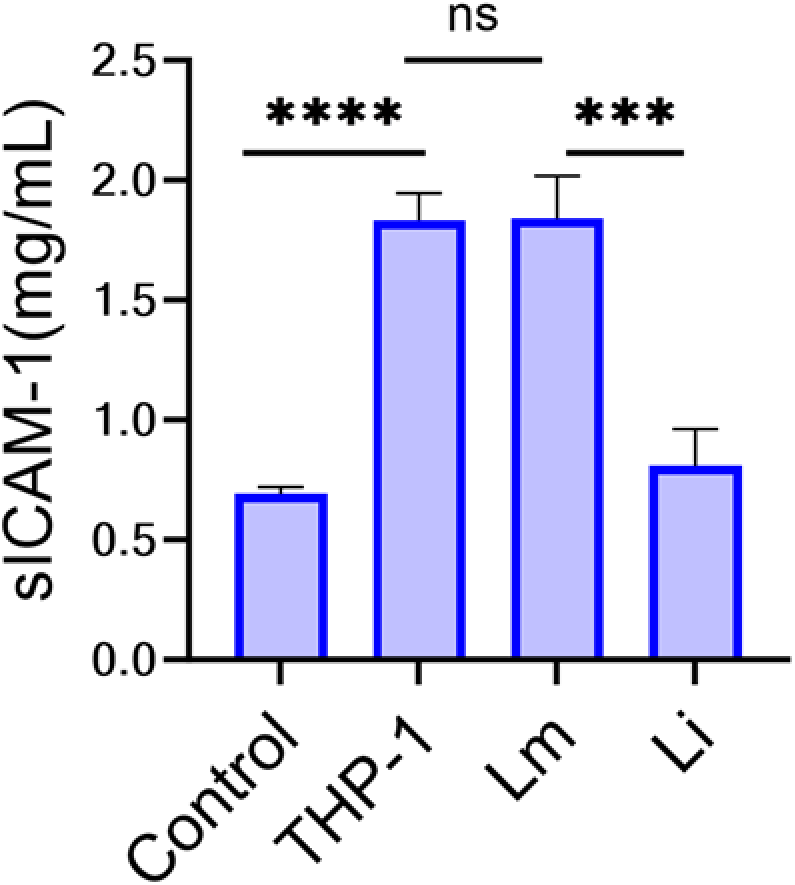
ELISA quantification of soluble ICAM-1 in apical supernatants. THP-1 monocytes induced an approximately two-fold increase in ICAM-1 secretion compared with no monocytes control. Data are represented as mean ± SEM from four independent experiments (n = 4 per treatment). Statistical significance was determined by one-way ANOVA with Tukey’s multiple-comparisons test (*, P < 0.05; **, P < 0.01; ***, P < 0.001).

**Table S1.** Adhesion molecule array identifying *Lm*–selective and enhanced endothelial factors associated with monocyte transmigration. †.

| **Molecule** | **Function** | **Fold Change (*Lm*)** | **Fold Change (*Li*)** | **Selectivity** |
| --- | --- | --- | --- | --- |
| **P-Selectin** | Leukocyte rolling/tethering | 2.25 | 1.54 | *Lm*-selective |
| **ICAM2** | Firm adhesion & transendothelial migration | 1.54 | 0.58 (↓) | *Lm*-induced; *Li* suppressive |
| **BCAM** | Monocyte-endothelial adhesion | 2.38 | 1.18 | *Lm*-selective |
| **VCAM-1** | Adhesion, transmigration signaling | 1.73 | 1.35 | Preferentially enhanced in *Lm* |
| **ICAM3** | Monocyte activation and adhesion | 2.25 | 2.53 | Shared response |
| **PECAM-1** | Diapedesis (transendothelial migration) | 1.99 | 2.02 | Shared response |
| **NCAM-1** | Junctional remodeling, immune trafficking | 3.81 | 3.70 | Shared, robust induction |
| **NrCAM** | Barrier modulation, immune interaction | 3.51 | 2.75 | Preferentially enhanced in *Lm* |

† Fold changes were calculated relative to untreated co-culture controls and normalized to internal positive controls; values represent the mean of pooled apical supernatants from four independent experiments analyzed in technical duplicate.

**Table S2:**
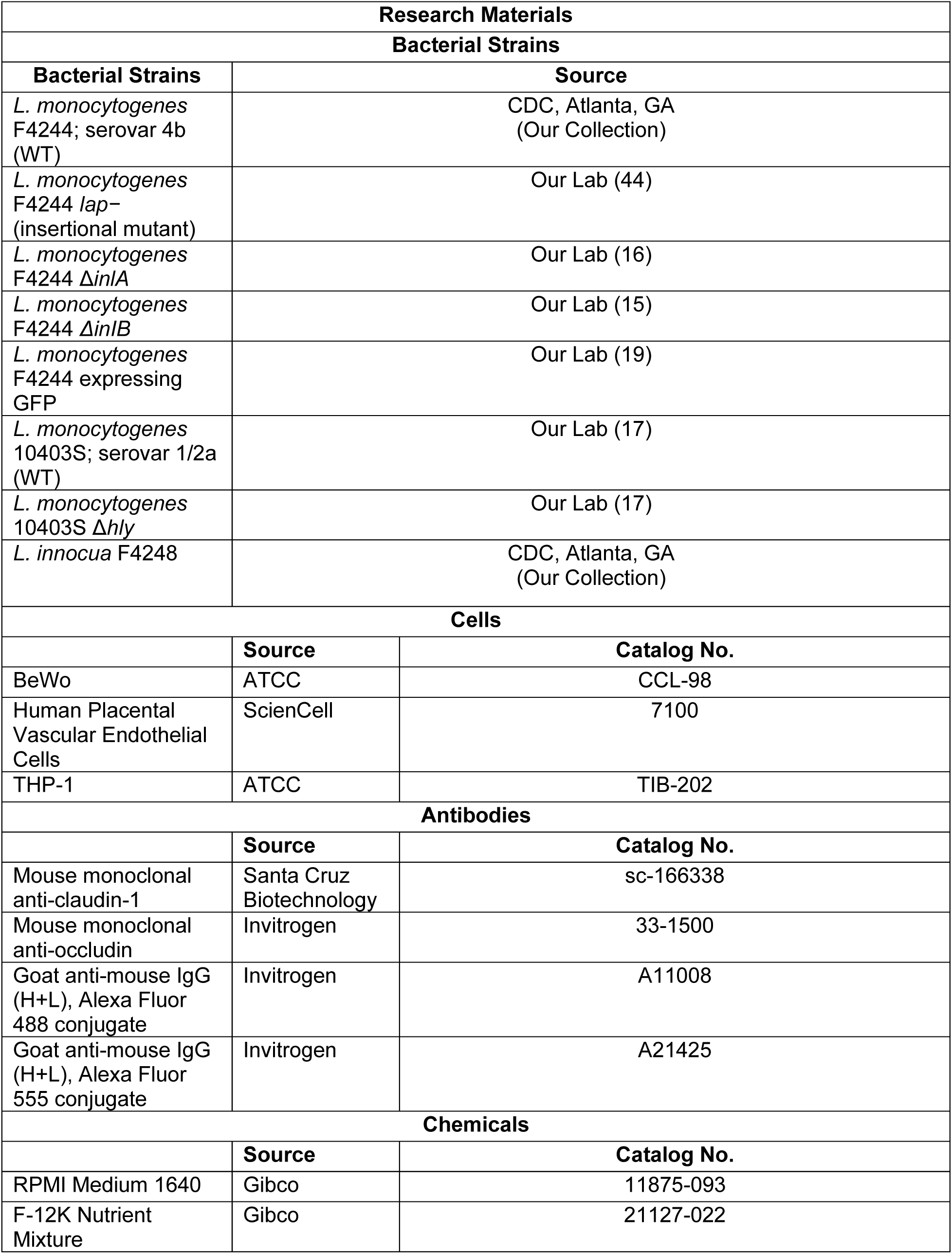
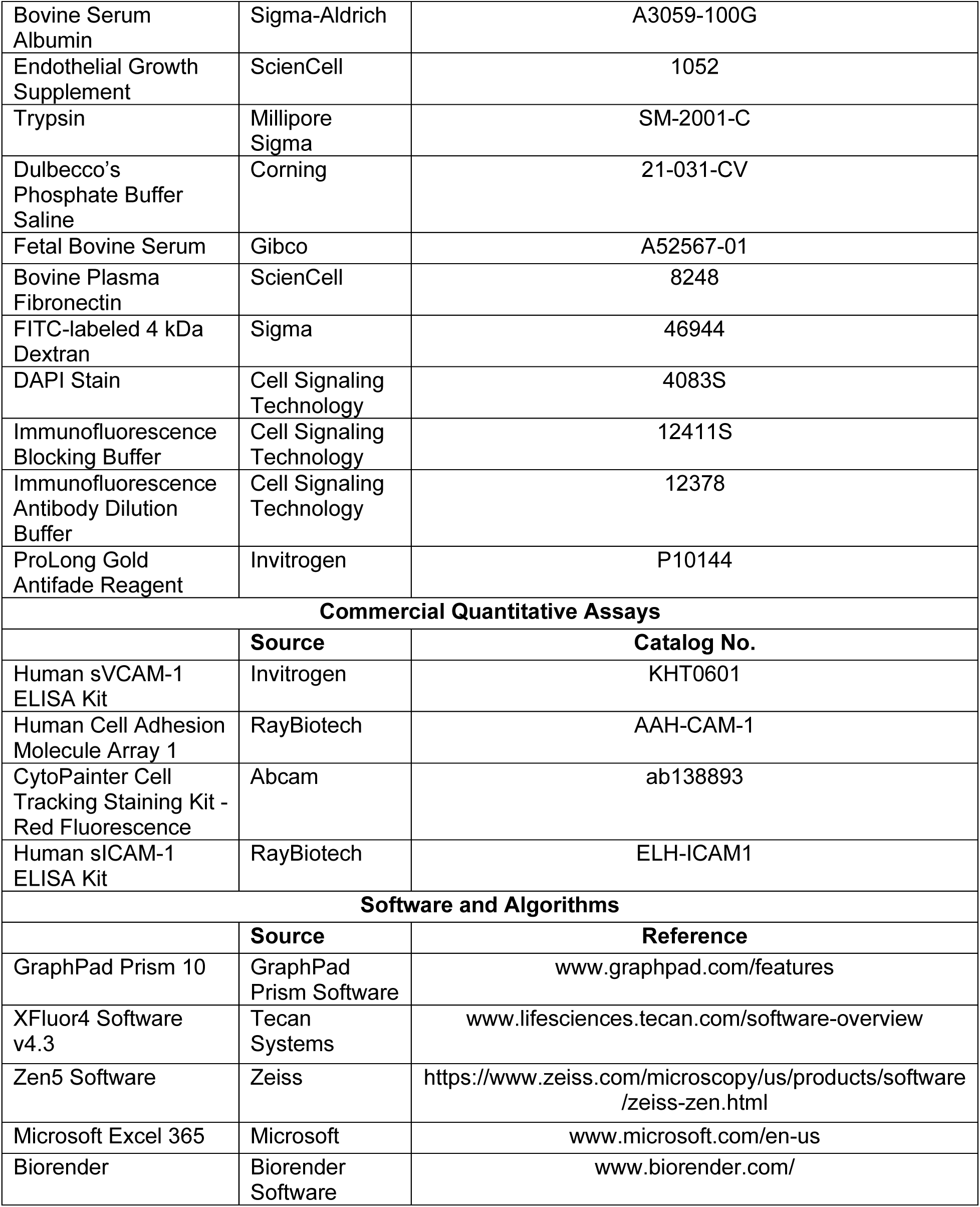
Key Resource Table of Materials Used.

